# Biosystematic Studies on the Status of *Solanum chilense* (Dunal) Reiche

**DOI:** 10.1101/2020.06.14.151423

**Authors:** Andrew R. Raduski, Boris Igić

## Abstract

- Members of *Solanum* sect. *Lycopersicum* are commonly used as a source of exotic germplasm for improvement of the cultivated tomato, and are increasingly employed in basic research. Although it experienced significant early and ongoing work, the taxonomic status of many wild species in this section has undergone a number of significant revisions, and remains uncertain.
- Here, we examine the taxonomic status of obligately outcrossing Chilean wild tomato (*Solanum chilense*) using reduced-representation sequencing (RAD-seq), a range of phylogenetic and population genetic analyses, crossing data, and morphological data.
- Overall, each of our analyses provides some weight of evidence that the Pacific coastal populations and Andean inland populations of the currently described *S. chilense* represent separately evolving populations.
- Despite its vast economic importance, *Solanum* sect. *Lycopersicon* still exhibits considerable taxonomic instability. A pattern of under-recognition of outcrossing species may be common across flowering plants. We discuss the possible causes and implications of this observation, with a focus on macroevolutionary inference.

## Introduction

The tomato and its close relatives (*Solanum* sect. *Lycopersicon*) form a small group with 13 currently recognized species (Peralta et al. 2008). The section is of exceptional interest because it harbors one of the most important commercial crops as well as its closely related wild species, which comprise an important germplasm resource for agronomic cultivar improvement (Rick and Chetelat 1995). The group also played a considerable role in foundational research on plant breeding systems, genetics, and the evolution of plant species (Eshed and Zamir 1995; Frary et al. 2000; Moyle and Nakazato 2010). The seminal taxonomic work by Rick and Lamm (1955) reflected a strong influence of Dobzhansky (1937), who defined species as “groups of populations, the gene exchange between which is limited or prevented in nature by one, or a combination of several, reproductive isolating mechanisms.” Rick and Lamm (1955) proposed that, in addition to accumulation of reproductive isolation, groups of populations may amass differences in other traits and “reasonably deserve recognition as separate species.”

Despite significant early work, the taxonomic status of tomato species has experienced a number of significant revisions (Peralta et al. 2008). Notably, one of the most genetically and phenotypically diverse species (*S. peruvianum* s.l.) now consists of four separate species. Species delimitation among members of the section is made difficult by a number of factors, including relatively stable population sizes, geographically widespread occurrence, recent divergence, a long history of obligate outcrossing enforced by self-incompatibility (SI), and possible introgression (Graham 2005; Pease et al. 2016; Böndel et al. 2015; Beddows et al. 2017). The motivation for the present work is related to the general reasons for taxonomic instability of the group, but has a relatively unusual origin.

Preliminary findings in the course of work on the evolution of self-incompatibility locus (S-locus) in *Solanum chilense* (Dun.) Reiche (Igic et al. 2007), uncovered a pattern of differentiation within functional alleles at the S-locus, pointing to possible substantial divergence between populations. Such differentiation is not likely within panmictic species, in part because all gene copies within the same functional allelic class are expected to coalesce ca. 10^2^ − 10^3^ generations, far faster than gene copies at a neutral locus, expected to coalesce ca. 10^6^ − 10^7^ generations (Vekemans and Slatkin 1994). We therefore reasoned that within-S-allele variation may provide a sensitive marker indicating that these populations may be so diverged as to represent cryptic species. It would not be peculiar to imagine that this scenario is common: SI species, which tend to have vast effective population sizes (Pease et al. 2016), could produce confusing patterns of morphological variation, and render difficult the ordinary search for a diagnostic taxonomic character. As a result, we initiated a research program aimed at examining the patterns of ecological, geographical, morphological, genetic, and crossing relationships among the populations of *S. chilense*, one of the most polymorphic wild tomato species, which is generally self-incompatible and spans particularly unlikely geographic and altitudinal ranges, with populations separated by the driest desert on the planet (Chetelat et al. 2009). Taken together with previous data (Graham 2005; Böndel et al. 2015; Beddows et al. 2017) these observations strongly hint that the current description of *S. chilense* demands closer examination.

The definition of species and inference of their boundaries and numbers continues to confound generations of systematists and evolutionary biologists. Diagnosable discontinuities between and among populations and species can arise as a result of common, continuous processes, and it is often unclear that steady accumulation of sequence data can provide clear guidance or resolve long-standing disputes. For example, local adaptation and population structure across plant populations are problematic in this sense, because they can create patterns that are not easily differentiated from those that appear as outcomes of interspecific divergence (Jordan 1846; Clausen et al. 1939, 1951) under any given conceptualization of species.

Despite some notable philosophical progress in attempts at a unified framework, vastly enhanced scale of genetic data and corresponding statistical approaches, species delimitation is limited by accumulation of convincing ecological and phenotypic data (De Queiroz 2007; Sobel et al. 2010; Sukumaran and Knowles 2017). A key problem is that different approaches to species delimitation can yield non-randomly varying conclusions regarding the boundaries (and numbers) of species, both relevant in the context of many other biological questions, including macroevolutionary inference. Myriad described problems plague approaches to macroevolutionary questions (Goldberg and Igić 2008; Maddison and FitzJohn 2015; Louca and Pennell 2020), but one under-appreciated confounding aspect concerns the species problem. For instance, we expect that taxonomic ‘splitting’ may associated with self-fertilizing taxa, because local inbreeding results in many diagnosably (morphologically) distinct entities, whereas ‘clumping’ may be associated with outcrossing, because it can cause high variation and few diagnosable characters. Such association of any character that scales effective population size potentially confounds inference of speciation and extinction rates, because species delimitation practices can mimic speciation and extinction.

Here, we apply the classic biosystematic approaches in order to ascertain the taxonomic status of an obligately outcrossing *S. chilense* and compare the results to similar studies in the related members of *Solanum* sect. *Lycopersicon*. Specifically, we (1) present eco-geographic variation with occurrence data and species distribution modeling, (2) perform controlled crosses and estimate the strength of early-acting post-mating barriers, (3) examine the distribution of morphological variation, and (4) establish phylogenetic and genetic relationships within *S. chilense*, using genome-wide restriction-site-associated DNA sequencing (RADseq), coalescent-based species tree estimation methods, and non-hierarchical methods. Each of the analyses shows a common discontinuity, consistent with the presence of at least two young separately evolving lineages within the presently described taxon.

## Materials and Methods

### Study Organism

*Solanum chilense* (Dunal) Reiche are herbaceous perennials with a woody base erect or decumbent growth, 0.5-1.5m tall and 1m wide when found isolated along cliff- and roadsides, but commonly spreading to 2-4m (or more) inside deep Andean volcanic canyons carved by snowmelt and sparse rain. Coastal natural populations instead occupy wide canyons below infrequent mist-capturing hill oases called “lomas formations” (Dillon 2005), scattered among the Chilean coastal hills (0-1000m) running along the Pacific Ocean, approximately 600km from Taltal to south of Iquique. Plants tend to inhabit wide sloping canyons with apparent subsurface flow and artificially formed water channels along roadsides, and in this region they are generally much shorter and decumbent. Inland populations stretch along the western slopes of the Andean cordillera (2000-3600m), from the vicinity of Salar de Atacama northward, well into central Peru (800-2000m). These montane populations are densest, featuring the largest plants, which generally occupy low rocky hillsides and dry beds with evident subsurface flow in the northern part of the range, as well as spectacular Andean slot canyons at the southern inland range margin.

Intriguingly, these discontinuous coastal populations and Andean populations are not connected by a common watershed in this area. In the southern portion of the range, Andean populations are very close to coastal populations (100-200km), despite being strictly separated by a hyper-arid buffer zone (Fig. 1). Andean populations of *S. chilense* represent the sampled altitudinal maxima, and coastal populations the sampled southern range limit, of all species in *Solanum* section *Lycopersicon* (Chetelat et al. 2009).

Taxonomy of *S. chilense* has undergone numerous changes (Rick and Lamm 1955; Peralta et al. 2008). The original description of *Lycopersicon chilense* Dunal relied on the earliest known collection of *S. chilense*, sampled by Gaudichaud in 1839 (Rick and Lamm 1955), from a coastal population at Cobija, Bolivia. Philippi (1860) cited both inland and coastal locations in a description of a new species, *Lycopersicon atacamense*, however both accessions were later included in the polymorphic *S. peruvianum* species complex. The current species status of *S. chilense* was determined based on studies of crossing relationships with other tomato species, somewhat distinct morphology, and a geographic range that is largely allopatric or parapatric with respect to other species formerly in the *S. peruvianum* complex (Rick and Lamm 1955). Reportedly distinguishing characteristics of *S. chilense* include structure of sympodial units, overall plant length, the ratio of peduncle length to inflorescence branch length, qualitative assessment of pubescence, stem color, staminal column curvature, as well as geography (Rick and Lamm 1955; Peralta et al. 2008). And yet, none of these are strictly diagnostic, as each is known to vary dramatically between and/or within populations (Peralta et al. 2008; A.R.R. and B.I., unpub. data). Our sampling scheme was informed by previous work and broadly aimed to discern relationships within and among the Chilean populations of *S. chilense*, as well as their geographic delimitation (Graham 2005; Igic et al. 2007).

**Figure 1:**
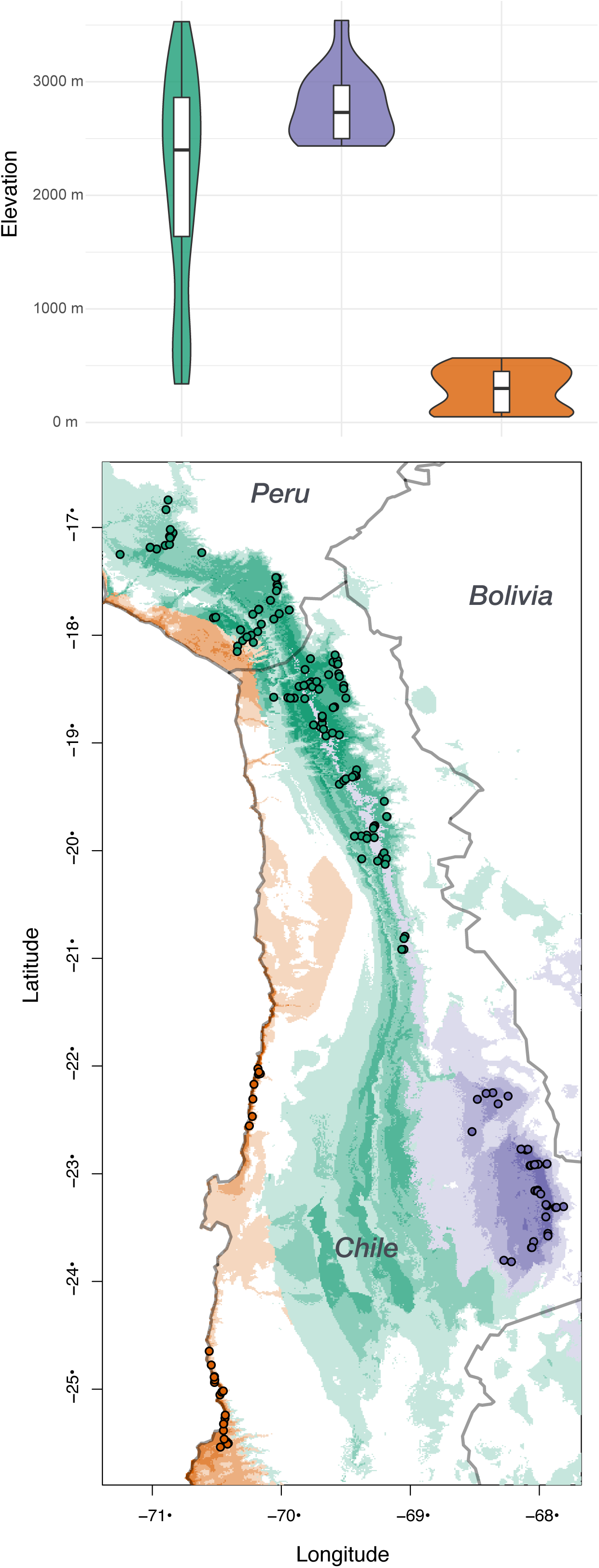
Geographic distribution of *Solanum chilense*. The top panel shows elevational distribution of the studied, binned and colored by regional designation: green–northern inland populations, purple–southern inland populations, and orange–southern coastal populations. The bottom panel shows occurrence and sampling locations, along with a range projected by species distribution models for occurrences from each of the three *a priori* designated population groups. Note that the large swaths of projected ranges in the middle of the map (without any samples), contain unsuitable habitat where superficially appropriate relief exists, but lacks Andean or lomas formation water sources. See text for details.

### Species Distribution Modeling

We constructed population distribution models based on climatic variables for each of three *a priori* determined groups, supported by phylogenetic and morphological analyses. We mainly employed the models for visual display and assessment of differences in distributions among the units in the *S. chilense* complex, because it appears that absence of a key ecological variable associated with occurrence of this species group, availability of underground water sources, results in strongly inflated predicted ranges. This omission notwithstanding, along with similar previous studies (Nakazato et al. 2008, 2010), our predictions provide an improvement over hand-colored maps (e.g. Igic et al. 2007).

Initially, we obtained geo-referenced presence data from the Global Biodiversity Information Facility (GBIF) database, but found them virtually unusable after careful curation, on account of many replicated and propagated errors (beyond the commonly encountered errors in such databases), including wildly incorrect locations. We list examples here, because we are unaware of published disclaimers concerning this taxon. In one instance (Gaudichaud’s original collection; see above) the centroid of Bolivia is given for the geolocation, on account of the original collection stating the country name. But it is well-documented that this sample is from the vicinity of Cobija, claimed by Chile since the War of the Pacific a century ago, but the database entry failed to take notice. (In fairness, this location is also claimed by Bolivia; nevertheless that location is off by several hundred km) Some locality data appear to have been entered with rounded (to nearest degree integer) low precision, or with incompatible listing of altitude and geolocation. Additionally, many data points trace to the Charles M. Rick Tomato Genetics Resource Center (TGRC) collections, which subsequently amended the originally listed coordinates. We therefore omitted GBIF for occurrence data used in species distribution models of this particular species group.

For the final georeferenced data collection, used in species distribution modeling, we instead relied on: (1) our own observations collected over three field trips December 1993 to April 1994 (W.A. Smith), April 2007 (B.I.), and January 2011 to May 2012 (A.R. & B.I.), (2) observations databased by the TGRC (Chetelat et al. 2009), (3) Georeferenced data found in the primary literature often not specifically concerned with this taxon, and (4) Georeferenced metadata from photographs, found by examining crowd-sourced websites. The occurrence data for this taxon are included in the Supplementary Materials.

We employed the processed occurrence data to fit and evaluate a maximum entropy species distribution model for each population, using a MaxEnt software implementation in the R-dismo package (Phillips et al. 2006; Hijmans et al. 2017) for R v. 3.4.1 (R Core Team 2017). Pseudo-absences were generated from 1000 point samples within a range restricted to the relatively narrow known occurrences range. We divided the species presence data into three equal partitions and used k-fold cross-validation. Two of these partitions were used to train the models with 30-arcsecond resolution climate data from 10 of 19 bioclim layers as predictors (Hijmans et al. 2004), after removing ones with significant correlations. We calculated performance by cross-validation, using Area Under the Receiver Operator Curve (AUC) statistic (Hijmans et al. 2017). The AUC is expected to vary from 0.5—for a model that performs no better than random—to 1.0 for perfect ability to predict presence versus absence. Background points were sampled from a rectangular extent, bounded by longitudes 75 and 67 (West), and latitudes 46 and 41.5 (South).

We used the probability surfaces of niche models obtained from MaxEnt to calculate the *I* -statistic of Warren et al. (2010). We estimated *I* for each pairwise set of populations using the function `nicheOverlap` implemented in R-dismo. The statistic is based on Hellinger Distances, and ranges from 0 (no overlap) to 1 (the distributions are identical). For clarity of visual display, we discretized SDM predictions by using areas with the occurrence probability cut-off at 0.80. We clipped the displayed map in Fig. 1 to highlight the regions of observed sympatry or parapatry, and larger maps (not shown) do not contain areas of significant overlaps.

### Genetic Differentiation Analyses

#### Population sampling

We sampled individuals from seventeen populations spanning the species range but focused on Chilean populations (Igic et al. 2007,Table 1). The sampling effort focused on three geographic areas corresponding to previously uncovered genetic groups within the species (Graham 2005; Igic et al. 2007): northern inland Chile (8 populations, 19 individuals), southern inland Chile (3 populations, 11 individuals), and coastal Chile (4 populations, 22 individuals). Additionally, four *S. chilense* individuals from two central Peru populations and eighteen *S. peruvianum sensu stricto* individuals from seven populations throughout the southern part of its species range were also sampled to help contrast intraspecific differentiation with interspecific divergence. Voucher specimens for the principal starting material, not originating from TGRC were deposited at the Missouri Botanical Garden (collector 37947, W. Allen Smith; collections MOBOT-WAS14 to -WAS37). We amended this sampling effort with seeds obtained from the UC-Davis C.M. Rick Tomato Genome Research Center (TGRC), which generously provided seeds of Peruvian *S. chilense* accessions (LA1917, LA1938, and LA0458) and Peruvian *S. peruvianum sensu stricto* accessions (LA0372, LA1977, LA2732, LA4318, LA4325). We also ordered one TGRC accession of *S. pimpinellifolium* (LA1245), which was included as an outgroup in later analyses.

**Table 1.** Sampling locality information for populations and individuals used throughout study. Geographic area codes are listed after population name (*S. chilense*: E = central Peru, N = northern inland range, S = southern inland range, C = southern coastal. *S. peruvianum*: PN = northern range samples, PS = southern range samples). Asterisks in the crossing, data morphology, or genetic columns signify inclusion of the population in the respective dataset. Number of samples and accession IDs refer to samples in phylogenetic study.

| Population | Data |  |  |  |  |  | Sequencing<br>Accessions |
| --- | --- | --- | --- | --- | --- | --- | --- |
|  | Latitude<br>°S | Longitude<br>°W | Crossing | Morphology | Genetic | No.<br>Samples |  |
| *S. chilense* |  |  |  |  |  |  |  |
| Llauta (E) | -14.27 | -74.92 |  |  | * | 2 | LA1917-01, LA1917-02 |
| Salsipuedes (E) | -15.68 | -73.83 |  |  | * | 2 | LA1938-01, LA1938-02 |
| Tacna (N) | -17.97 | -70.18 |  |  | * | 2 | LA0458-01, LA0458-02 |
| Socoroma (N) | -18.16 | -69.57 | * |  | * | 1 | WAS365-01 |
| Livilcar (N) | - 18.3 | -69.43 | * |  | * | 1 | WAS237-02 |
| Azapa (N) | -18.41 | -69.97 |  | * | * | 3 | BI786F, BI786G, BI786J |
| Cardones (N) | -18.46 | -69.82 |  | * | * | 6 | BI766, BI770, BI775 |
|  |  |  |  |  |  |  | BI776, BI777, BI779 |
| Timar (N) | -18.45 | -69.41 | * |  | * | 5 | WAS23-01, BI791H, BI791J<br>791P, 791R |
| Mocha (N) | -19.79 | -69.29 | * |  | * | 1 | WAS384-01 |
| Toconce (S) | -22.25 | -68.42 | * |  | * | 1 | WAS436-03 |
| Ruta 27 (S) | -22.92 | -68.04 | * | * | * | 9 | BI502, BI517, BI522, BI523, BI548<br>BI554, BI558, BI569, WAS462-03 |
| Talabre (S) | -23.33 | -68.00 | * |  | * | 1 | WAS85-02 |
| Tocopilla (C) | -22.08 | -70.18 |  | * | * | 5 | BI737J, BI737N, BI737R<br>BI737U, BI737W |
| Paposo (C) | -24.88 | -70.52 |  | * | * | 5 | BI736D, BI736L, BI736M<br>BI736P, BI736Q |
| Caleta Agua Buena (C) | -25.24 | -70.43 |  | * | * | 6 | BI705, BI707, BI722<br>BI728, BI729, BI732 |
| Taltal (C) | -25.48 | -70.43 | * | * | * | 6 | BI700A, BI700B, BI700D<br>BI700F, WAS44-01, WAS51-02 |
| *S. peruvianum* |  |  |  |  |  |  |  |
| Ancash (PN) | -9.94 | -78.23 |  |  | * | 2 | LA0372-01, LA0372-02 |
| Oucocto (PN) | -12.12 | -76.45 |  |  | * | 1 | LA1977 |
| Moquella (PS) | -19.40 | -69.60 |  |  | * | 2 | LA2732-01, LA2732-02 |
| Sora-Molinos (PS) | -18.35 | -69.88 |  |  | * | 2 | LA4318-01, LA4318-02 |
| Caleta Vitor (PS) | -18.75 | -70.33 |  | * | * | 2 | LA4325-01, LA4325-02 |
| Azapa (PS) | -18.58 | -70.03 |  | * | * | 6 | BI788H2, BI788K, BI788N<br>BI789H, BI789K, BI789N |
| Poconchile (PS) | -18.58 | -69.94 |  | * | * | 3 | BI790C, BI790E, BI790J |

#### RADseq library preparation, sequencing, and data assembly

We used fresh or silica-dried leaf tissues for DNA extraction with the CTAB method (Murray and Thompson 1980). Quality and quantity of extracted DNA were measured with a NanoDrop2000 and Qubit. Restriction-site-associated DNA sequence (RADseq) libraries containing uniquely barcoded DNA samples were created using a single *PstI* enzymatic digestion of sample DNA following the protocol of Elshire et al. (2011). We checked the quality of individual RADseq samples on an Agilent 4200 Tape Station system and samples were combined in equimolar concentrations into sequencing libraries at the the DNA Services facility within the Research Resources Center at the University of Illinois at Chicago. Libraries were then sequenced with Illumina HiSeq2500 technology to generate 100bp paired-end reads at the Roy J. Carver Biotechnology Center at the University of Illinois.

We demultiplexed and trimmed adaptor and barcode sequences from raw reads using Sabre (https://github.com/najoshi/sabre). Reads were then cleaned using the `SLIDINGWINDOW` setting in Trimmomatic (Bolger et al. 2014) and clipped when the average Phred score fell below 15 within a four base pair sliding window. The resulting reads were then mapped to the *S. lycopersicum* genome, build SL2.50 (Fernandez-Pozo et al. 2015), using the default settings of the MEM algorithm of BWA (Li and Durbin 2009). Sequence reads from all samples were grouped by mapped genomic location coordinates using the `pstacks` module of *Stacks* (Catchen et al. 2013). Sequences from RADseq loci found in at least 90% of sampled individuals were retained for phylogenetic analyses. The same set of loci were used for population genetic analyses, however we focused only on populations with dense sampling. We also analyzed alternative datasets, containing loci found in varying numbers of individuals, but the results do not qualitatively differ from those presented.

#### Phylogenetic relationships

Phylogenetic relationships across samples were estimated using two methods. First, we used an intuitive, simple, and convenient approach: concatenation with tree-based inference. RAxML v8.2.8 (Stamatakis 2014) with a concatenated sequence alignment and GTR+G model of nucleotide evolution. Node support values were estimated by 1,000 bootstrap replicate searches from random starting phylogenies. Concatenation-based approaches have been the subject of some criticism (Ree and Hipp 2015), especially regarding statistical inconsistency and its failure to capture coalescent process of unlinked sites (Chifman and Kubatko 2014). Consequently, we approach these results with some skepticism, but its supposedly rampant failures are far from certain, and a number of potentially misleading results are thought to be reasonably easily uncovered (Ree and Hipp 2015).

Nevertheless, we are interested in a statistically consistent species tree, so we additionally implemented SVDquartets, a coalescent-based tree inference method (Chifman and Kubatko 2014). Briefly, this method calculates optimal quartets (unrooted four-taxon relationships) for unlinked nucleotide site patterns under the coalescent. All quartet trees are then summarized and assembled to estimate relationships among samples. We examined all possible quartets among our samples, making no prior assumptions about population clustering, and estimated node support using 1,000 bootstrap replicates in PAUP* v4.0a150 (Swofford 2003). SVDquartets is thought to be reasonably robust to violations of its assumptions, including linkage, as well as missing data.

#### Population clustering with Structure

Next, we estimated the presence of distinct populations, assign individuals to populations, and identify admixed individuals within *S. chilense*, using the model-based algorithm implemented in Structure (Pritchard et al. 2000; Falush et al. 2003). Each Structure run infers the proportion of ancestry of each individual assigned to one of a predefined number of genetic clusters (K). Fifteen replicate analyses were performed across a range of K values (K = 1 – 12). Our Structure input data was restricted to a single random SNP from each RADseq locus as implemented in *Stacks*. Each analysis contained 100,000 burn-in iterations, followed by 100,000 data-collecting iterations of the admixture model with uncorrelated allele frequencies. We chose the optimal number of clusters as the K with the highest ΔK, a robust selection criterion for identifying correct higher level structuring of populations based on changes in likelihood surfaces with increasing values of K, in Structure Harvester (Evanno et al. 2005; Earl et al. 2012). Data from replicates were summarized using CLUMPP (Jakobsson and Rosenberg 2007) and visualized using Distruct (Rosenberg 2004). Genetic differentiation of population allele frequencies (Wright’s *F*_*st*_) was calculated between pairwise combinations of populations within and between *S. chilense* regions and *S. peruvianum* (Weir and Hill 2002). The average number of pairwise nucleotide differences per base pair were recorded as a measure of genetic variation. We separate pairwise comparisons into within (*π*) and between population (*D*_*xy*_) to better understand how variation is distributed. The genetic structuring of *S. chilense* populations were also investigated with a principal components analysis (PCA) of genotypes at a subset of loci filtered for linkage disequilibrium. The coordinates of the first two principal component axes of each sample were Procrustes transformed onto their geographic coordinates to better illustrate the relationship between genetic variation and geography. While the interpretation of Structure analyses and *F*_*st*_ values can be problematic when populations are not discrete (individuals are arrayed along a geographic continuum) we are reasonably confident that this is not the case for *S. chilense*, because the present and recent climactic features of Atacama desert show stark discontinuities and enable visual confirmation of absence across the vast barren landscape.

#### S-allele haplotypes

We previously sequenced and verified allelic identity of a large number of S-alleles from *Solanum chilense* (Igic et al. 2007). In the course of that study, we uncovered a number of single feature polymorphisms within allele classes, which could uniquely identify the coastal populations from the rest of the described taxon. Such divergences are unlikely in panmictic populations or in case of inter-species introgression (Castric et al. 2009). Consequently, we sought and found representatives of a single allele class in multiple populations and sequenced them in order to characterize within-allele polymorphism at the S-locus.

Briefly, we extracted DNA from leaves as described for RADseq (below), and total RNA from styles, followed by cDNA synthesis, primed amplification, and Sanger sequencing as described in Igic et al. (2007). We found several polymorphic alleles, but successfully recovered only two (SX-30 and S3-6) in all sampled populations (coastal, south and north inland). This is unsurprising, because 35-45 alleles segregate in populations of this species at low frequencies, and very large sample sizes are required to find the same allele(s) in multiple populations. The accessions with variants of S3-6 (corresponding to GenBank accession EF680086.1 sequence, Igic et al. 2007) were: (1) coastal populations: WAS42-2, WAS47-2, WAS44-3, (2) inland north: WAS23-1, (3) inland south: WAS462-3, and one outgroup accession TGRC, (4) *Solanum peruvianum* LA1556 (individual 08B87) The accessions with variants of SX-30 (corresponding to GenBank accession EF680100.1 sequence) were: (1) coastal populations: WAS44-3, WAS50-3, WAS53-5, (2) inland north: WAS23-1, (3) inland south: WAS462-3, and two outgroup accessions from TGRC, (4) *Solanum pennellii* LA1674 (individual 08B64), *Solanum arcanum* LA2152 (08B8). Haplotype networks shown in Fig. 2 were created using the function `pegas::haploNet()` (Paradis 2010). We will detail the remainder of this S-allele search in a separate publication, but we present an excerpt of the work here, because it is instructive in its independent confirmation of genetic divergence patterns.

**Figure 2:**
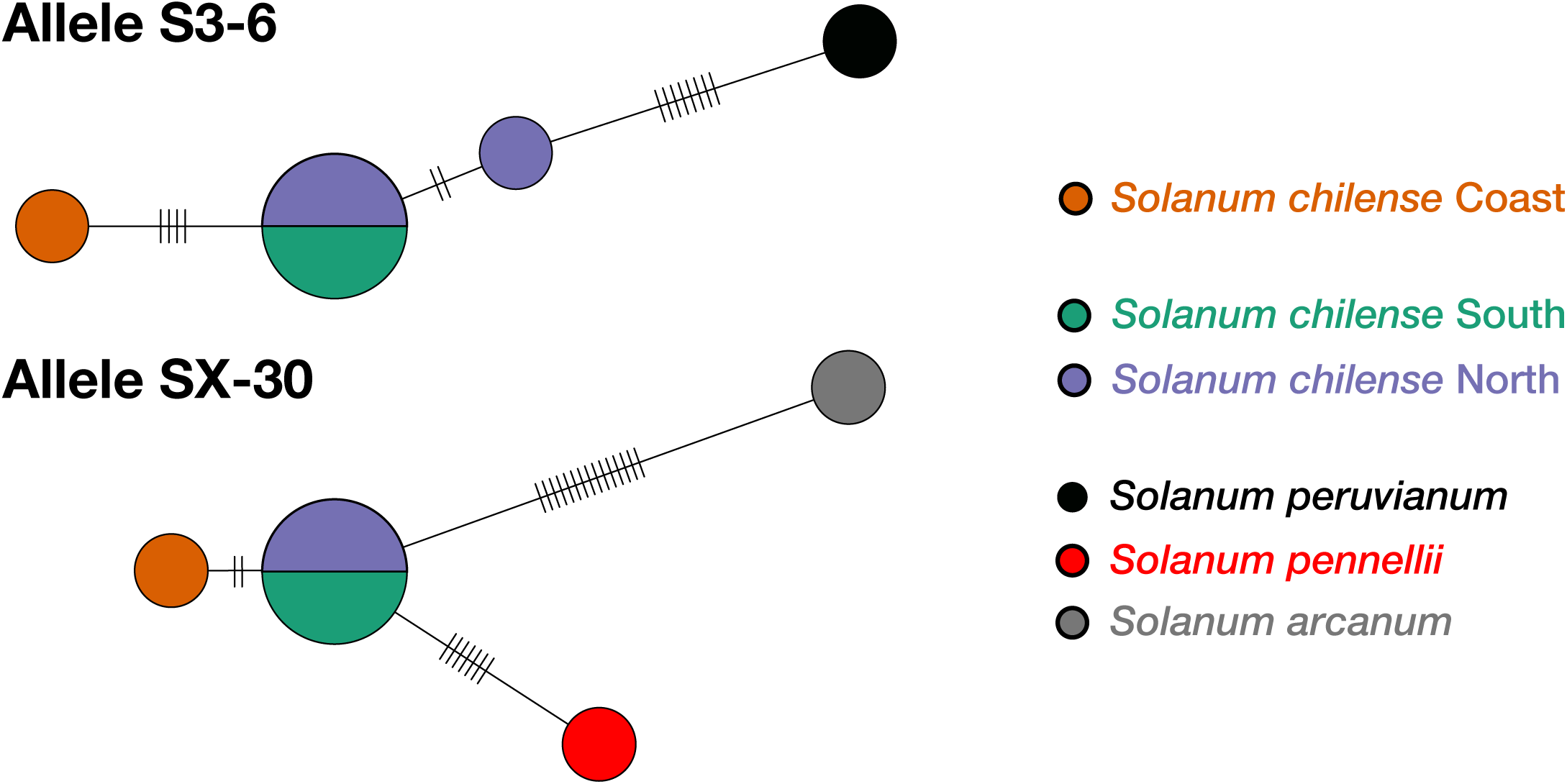
S-allele haplotype network for alleles S3-6 and SX-30. These two alleles are found in *Solanum chilense* from all three geographic regions we sampled here and previously (Igic et al. 2007), as well as in closely related species. S3-6 was sampled five times in *S. chilense* and once in *S. peruvianum*. SX-30 was sampled five times in *S. chilense*, and once apiece in *arcanum* and *S. pennelli*. S-alleles have small effective population sizes, compared to other nuclear loci—on the order of 2*N*_*e*_/50 for a population with 50 segregating alleles—and they experience strong frequency-dependent selection. Therefore, they can be sensitive indicators of gene flow (see text for detail; Vekemans and Slatkin 1994).

### Morphological Differentiation

We measured multiple morphological characters from inland and coastal *S. chilense* and *S. peruvianum* populations (Table 1), many of which were previously identified as being variable in the group (Graham 2005; Peralta et al. 2008). Floral and inflorescence characters were measured directly from plants in natural populations, with the aid of digital calipers. Ideally, 10 measurements were taken for each floral and inflorescence character on each plant. In rare cases, individuals displayed fewer than 10 mature flowers, in which case we measured all available flowers. We measured the number of major intra-inflorescence branching events, flower number, intra-branch length, peduncle length, pedicel length, flower length, anther length, anther width, sterile anther length, stigma exertion, sepal length, and petal length.

Leaf characters were measured from photographs of leafs taken in the field. Three measurements for each leaf character were recorded per individual (Table **??**). We measured total leaf length, number of lateral leaflets, number of interstitial leaflets, terminal leaflet length, and distal leaflet length. A PCA was performed on mean character values for each individual using the *prcomp* function in R (R Core Team 2017). To assess how well the morphological measurements predicted group membership, linear discriminant analysis (LDA) was performed using the MASS package in R (Venables and Ripley 2002). Our field based measurements are certainly influenced by environmental variation. Although we did not perform experiments necessary to estimate trait heritabilities (Falconer et al. 1960), maternal half-sib families from coastal and southern inland individuals are separated by region along the major axes of phenotypic variation while grown in a greenhouse (Fig. Supplement). Some of the characters measured have also been shown to be highly heritable in other tomato species (Chitwood et al. 2013).

### Post-pollination Reproductive Isolation

We estimated the strength of a subset of intrinsic post-mating reproductive barriers by performing controlled glasshouse experiments, examining seed set, and assessing a proxy of offspring fitness. The crossing design relied on preliminary data indicating that coastal populations were possibly distinct, despite being geographically close to southern Andean populations. We performed reciprocal inter- and intra-population crosses between northern Andean populations, southern Andean populations, and coastal populations. We used 10 plants from each population, but a full reciprocal design would have resulted in a prohibitive number of crosses, given the time and number of flowers available. Instead, each plant was randomly paired for two reciprocal intra-population crosses, as well as inter-population crosses with two plants from each of the other populations.

All plants used in the crossing experiment were grown in the glasshouse at the University of California—San Diego, from seeds originally collected in by W.A. Smith (Igic et al. 2007). The numbers of seeds per fruit in all reciprocal inter- and intra-population crosses were counted as a proxy of the strength of post-mating, pre-zygotic, barriers. Seed size and germination rate were also recorded as proxies of seed viability, at slightly later post-zygotic stages. We compared success across population crossing classes with one-way ANOVA and assigned classes with Tukey’s honest significant differences (HSD) test (Zar 1999), as implemented in the R `stats` package (Venables and Ripley 2002).

## Results

### Geographic Survey and Species Distribution Modeling

The distribution of populations in the *Solanum chilense* complex is highly discontinuous (Fig. 1). Our ground surveys, and those conducted by others (Chetelat et al. 2009), indicate that the stark discontinuities between the inland and coastal populations in northern Chile are not artifacts of collecting effort. We find that the commonly scored climate and geography data, such as those found in bioclim layers (Hijmans et al. 2004), do not cover distributions of canyons, which form the vast majority of suitable habitat for this taxon in the Atacama desert. Nevertheless, such data appear somewhat useful.

The distribution of this taxon, as well as its near relatives, closely follows the climate of the discrete Köppen classification for Cold desert climate (BWk; north and south inland populations). The coastal populations are, however, closer to the southern portion of BWh in Chile. They are not found in the vicinity of, or north of, Rio Loa valley, where lomas formations are absent and absolute desert lines the coast up to the 19th parallel (Dillon 2005). Moreover, species distribution models apparently performed well, with AUC>0.95 in each case, performing far better than null geographic models. The mean niche overlap across three cross-validation subsamples was highest for northern inland and southern inland populations (*I*_*NS*_ = 0.54), and substantially lower for the two pairwise comparisons involving the coastal populations (*I*_*NC*_ = 0.32, *I*_*SC*_ = 0.01). This is somewhat unsurprising, given the vast differences in altitudinal range (2000 m) between the densest populations in the Andean cordillera and the Pacific coastal hills that comprise lomas formations. Although bioclim variables contain rainfall data, they do not account for snowmelt-fed aquifers and other underground water availability, which are likely key in determining the distribution of this and related species. Much of the projected niche overlaps occur in such water-free areas (such as Cordillera Domeyko, ca. 24.5° S, 69° W), unsuitable for plant growth, and indeed nearly totally devoid of any biomass. Consequently, these projections are likely to vastly overestimate suitable niche overlap. And yet, some fairly fine-scale geographic features are readily discernible in our coarse projections. For example, the grand forked canyon of Rio Camarones on its approach to the Pacific Ocean (19° S, 70° W) is visible as possibly suitable for the coastal taxon, as are the areas slightly to the north. Both are highly reminiscent of the southern coastal population habitats and presently occupied by *S. peruvianum*. Finally, the northern-most disjunct populations in Peru, which we omitted in our distribution models, remain poorly sampled and generally understudied. It is unclear whether they should be included within any taxa within the *S. chilense* complex, another currently described taxon, or a heretofore undescribed group.

### Genetic Differentiation

#### RADseq data assembly

After demultiplexing and filtering reads for low quality sites, an average of 11.8 million reads per individual were used for downstream analyses. The MEM algorithm of BWA successfully mapped 86% and 73% of clean reads from *S. chilense* and *S. peruvianum* samples, respectively, onto the domesticated tomato genome. The `pstacks` module of *Stacks* identified an average of 3600 loci in each sample. Further filtering of the dataset to remove loci with low read or sample coverage resulted in 1302 and 833 loci for phylogenetic and population genetic analyses, respectively. Summaries of the mean numbers of sequence reads obtained, numbers of sequence reads successfully mapped, and total numbers of unique loci produced from *Stacks* for each *S. chilense* group are reported in Table **??**.

#### Phylogenetic relationships

A concatenated sequence matrix of 129,974bp and a SNP matrix of 954bp were used by RAxML and SVD Quartets, respectively, to estimate phylogenetic relationships. Both reconstruction methods gave qualitatively similar tree topologies concerning the relationship of coastal populations among other *S. chilense*. All coastal individuals formed a monophyletic clade with 100% bootstrap support in RAxML and SVD Quartets phylogenies (Figs. 3,4). The relationships among Andean plants differed in the results from the two reconstruction methods (Fig. S1). RAxML reconstructed a larger paraphyletic clade of northern and southern inland individuals as sister to the coastal clade with low bootstrap support (31%). One single northern inland plant (786G) was placed outside of the larger inland and coastal clade. SVD Quartets placed a highly supported clade of southern inland individuals as sister to the coastal clade with low bootstrap support (28%) (Fig. **??**). Most northern inland individuals were estimated to form a monophyletic clade. Both reconstruction methods placed paraphyletic central Peru populations outside of the larger monophyletic Chilean coastal and inland clades (Figs. 3, 4). *Solanum peruvianum* individuals were always estimated to form monophyletic groups sister to all *S. chilense* individuals, although bootstrap support varied depending on the method used. SVD Quartets estimated 100% bootstrap support for the *S. peruvianum* clade. RAxML estimated 55% bootstrap support for the *S. peruvianum* clade.

**Figure 3:**
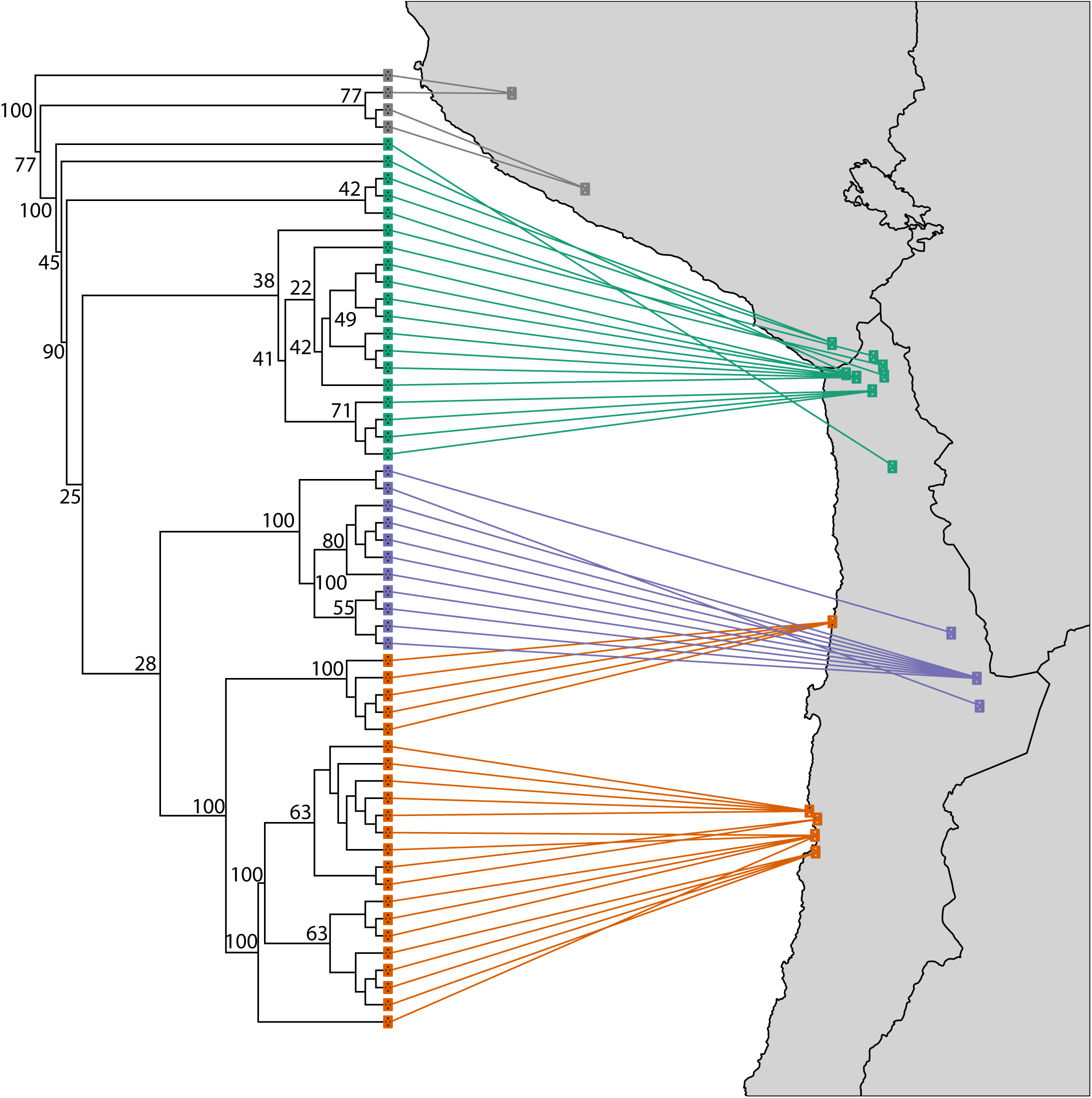
SVD Quartets phylogenetic reconstruction of *S*. chilense individuals. Node support was estimated using 1,000 bootstrap replicates. Support values are given on select nodes on the phylogeny. Northern inland (N; green), southern inland (S; blue), and coastal (C; orange) Chilean populations are shown, along with the far northern populations from central Peru (FN; grey). See Table 1 for detailed location information.

**Figure 4:**
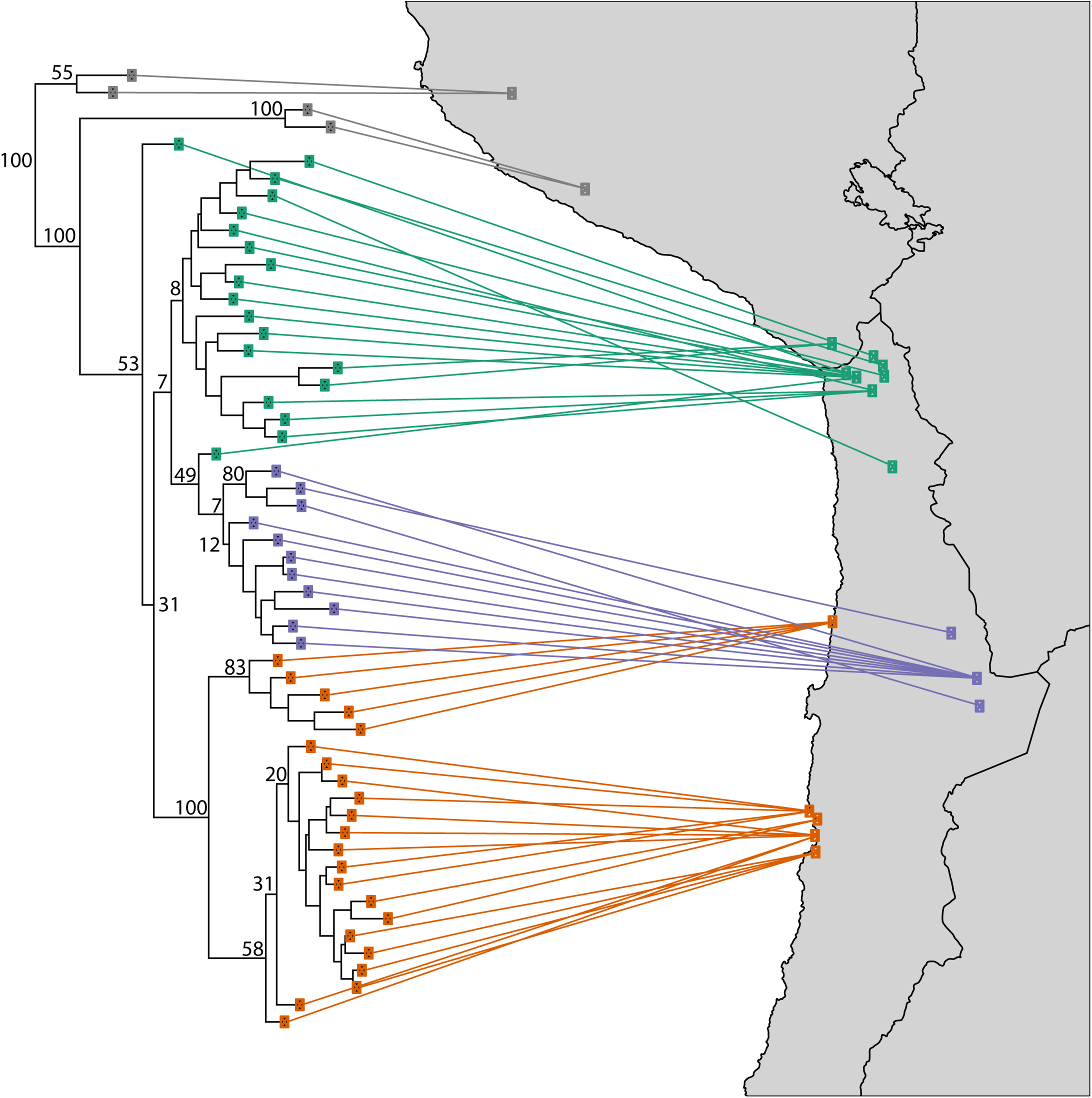
The maximum likelihood tree depicting phylogenetic relationships among the sampled *Solanum chilense* individuals, implemented in RAxML. The branch support values are given only on key nodes of the tree. They show the proportion of bootstrap trees that contain a given clade on the original ML tree, estimated using 1,000 bootstrap replicates. Northern inland (N; green), southern inland (S; blue), and coastal (C; orange) Chilean populations are shown, along far northern populations from central Peru (FN; grey). See Table 1 for detailed location information.

#### Population clustering with Structure

We identified K = 2 genetic clusters as the best supported model in our Structure analyses (Δ*K*_2_ = 440.44). The two inferred layers predominantly make up coastal *S. chilense* and inland Andean *S. chilense* ancestries. The analyses infer the majority of samples to contain ancestry from predominantly one genetic cluster. All but five samples contain at least 95% of ancestry from a single genetic cluster. The algorithm assigns three coastal samples from Tocopilla as containing admixed ancestry with <8.3% from the northern inland population, one coastal Caleta Aqua Buena sample 25.3% inland ancestry, and one northern inland sample from Socoroma containing 17.9% ancestry from the coastal genetic cluster.

Models with an additional cluster were the next best supported (Δ*K*_3_ = 99.93). The addition of an extra K did not qualitatively change the pattern of genetic structuring across samples. With K=3 no samples were inferred to contain more than 18.4% ancestry of the third additional genetic cluster (Fig. 5). All four central Peruvian *S. chilense* samples have 10–18.4% of their ancestry from the third genetic cluster and 81.4–90% shared ancestry with other inland samples. A single northern inland sample from Azapa contains 94.5% inland ancestry, 5.4% third cluster ancestry, and 0.1% coastal ancestry. The addition of a third genetic cluster increased the proportion of the minor ancestry group in coastal Tocopilla samples and the single coastal Caleta Aqua Buena sample with inferred mixed ancestry in the K=2 analysis. The estimated amount of inland ancestry in Tocopilla samples increased from 6.0% to 8.3% in the K=2 analysis to 19.0% to 27.7% in the K=3 analysis.

**Figure 5:**
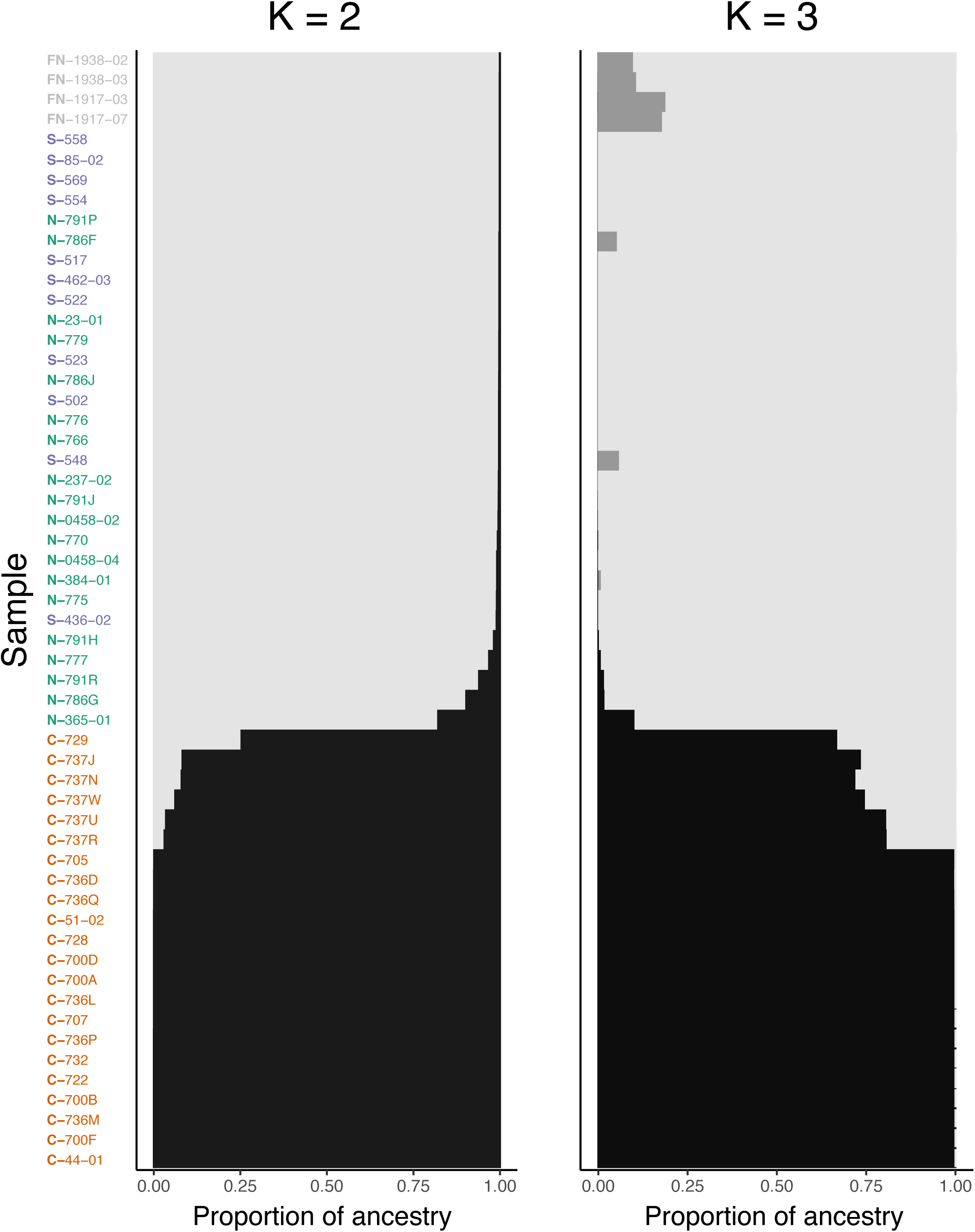
Estimated proportion of ancestry of *S. chilense* samples of K=2 (best supported) and K=3 clusters from Structure analyses. Coastal (C) vs. southern (S) and northern (N) inland populations are consistently placed into separate clusters.

Overall, levels of population allele frequency divergence were high across *S. chilense* populations (*F*_*st*_ ranged from 0.13 to 0.37) (Fig. S2). Within regions on average, coastal *S. chilense* populations were more diverged from each other (*F*_*st*_ = 0.20) than northern inland populations (*F*_*st*_ = 0.13). We were unable to calculate *F*_*st*_ within the southern inland region because only a single population was well sampled. Average between region divergence was lowest between southern and northern inland populations (*F*_*st*_ = 0.22). Coastal populations were slightly more diverged from southern inland (*F*_*st*_ = 0.37) than northern inland populations (*F*_*st*_ = 0.34). The levels of differentiation observed between coastal and both inland populations are near that of interspecific divergence between northern inland *S. chilense* populations and *S. peruvianum* (*F*_*st*_ = 0.39).

Our measure of genetic diversity, the average number of pairwise nucleotide differences per site, varied within (*π*) and between (*D*_*xy*_) populations across the landscape. We only interpret each result relative to comparisons within this study because estimating population genetic parameters from RADseq data is problematic (Arnold et al. 2013; Cariou et al. 2016). Within *S. chilense* the least genetic diversity is in coastal populations, however southern inland populations are comparable. Northern inland populations are the most diverse and have an average *π* value that is outside the range of coastal and southern inland *π*s.

Overall, levels of *D*_*xy*_ within *S. chilense* were comparable to *π* in northern inland populations. Comparisons between northern inland and coastal populations were, on average, higher than among inland populations suggesting that coastal populations have been longer isolated from the north. Most *D*_*xy*_ between coastal and southern inland populations falls outside of either region’s range of *π* measurements, indicating strong differentiation at the nucleotide level, perhaps due to reductions in local effective populations sizes.

#### S-allele haplotypes

We surveyed 96 individuals of *Solanum chilense* and recovered two S-alleles (S3-6 and SX-30) in all three sampled populations (coastal, south and north inland; Fig. 2). In each case, the S-alleles sequenced display variation consistent with considerable divergence between coastal and inland populations. We found three unique sequence variants of Allele S3-6. The first variant was found in one individual apiece from both inland populations. Another variant, two SNPs apart was found in another nearby northern inland population. Three individuals Coastal population each had the same (third) variant, four to six SNPs apart from the inland populations. We found two unique sequence variants of Allele SX-30. The first variant was found in two northern inland individuals and one southern inland individual. The second variant, two SNPs apart was found in all three sequenced coastal individuals. For visual comparison, Fig. 2) shows a haplotype network of intra-allelic sequence variants, likely functionally identical, from other species where they were found and sequenced.

### Morphological differentiation

The total morphological dataset contained 15626 floral or inflorescence and 2730 leaf measurements from 108 *S. chilense* plants and 1027 floral or inflorescence and 462 leaf measurements from 22 *S. peruvianum* plants. The first two principal components (PCs) explained 51.88% of variation within the morphological dataset (Table **??**). Leaf and inflorescence size variables had the largest loadings on the first PC. Leaflet size and leaf shape variables had the largest loadings on the second PC (Table **??**). Coordinates of the first two PCs of each sample and the mean coordinate values of samples from each group with ellipses encompassing 95% of the variation of *S. chilense* populations and *S. peruvianum* are plotted in Fig. 6. There is some overlap in the morphological PCA space of all groups, however the largest is observed between northern and southern inland *S*. chilense populations. A large portion of the coastal populations morphological variation is separated from that of inland populations along PC1. Morphological differences are also displayed by each group’s centroid coordinates in PC space. Coastal *S. chilense* populations are more separated from inland *S. chilense* populations, in Euclidean distance between centroid coordinates, than they are from *S. peruvianum* plants. The majority of individuals (90.86%) were assigned to their correct populations using linear discrimination analysis (Table **??**).

**Figure 6:**
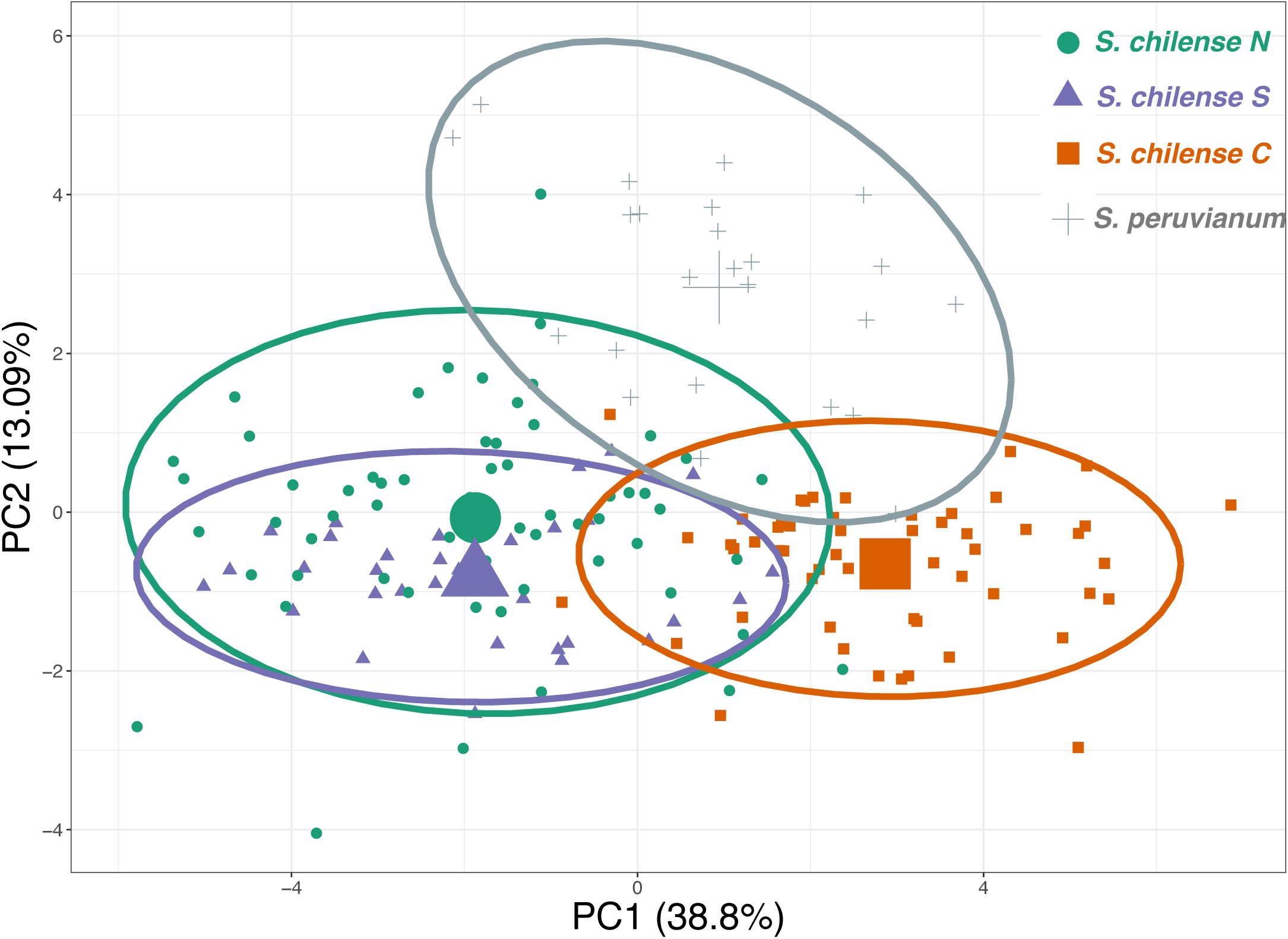
Individual and average group coordinates for the first two morphological principal components. Ellipses are drawn around 95% of the variation within each *S*. chilense geographic group and *S*. peruvianum (P). Large unfilled symbols represent mean coordinates for each geographic group.

### Reproductive Isolation within Solanum chilense

Intra and inter-population crosses (Fig. 7) produced a grand mean of 32.47 seeds, but overall crossing success was significantly different between cross types (*F* = 23.95, *df* = 8, *p* < 2 × 10^−16^). Inter-population crosses that included a coastal plant consistently produced significantly fewer seeds per fruit than all other cross classes. Four such cross types averaged 14.38 seeds per fruit, while inland inter-population crosses yielded an average of 42.25 seeds per fruit. Numbers of seeds produced from crosses within the coastal population were lower than other intra-population crosses, but not statistically significantly different from crosses between northern and southern inland plants.

**Figure 7:**
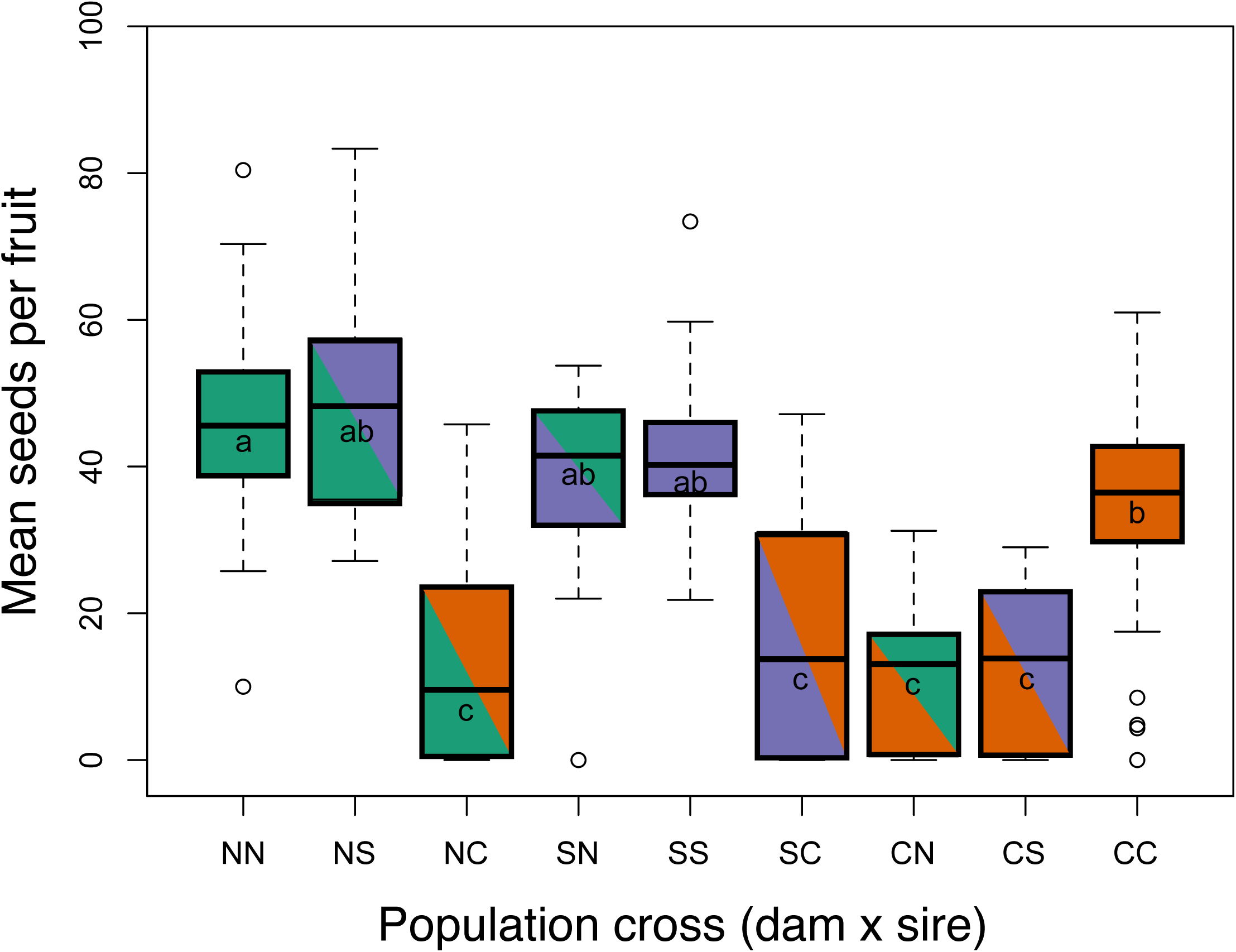
Post-pollination reproductive isolation between *S*. chilense populations. Each plot shows estimates for post-mating, pre-zygotic isolation, as seed set per fruit (233 crosses, 1046 fruits, 32,868 seeds total), grouped by population of origin of maternal and paternal parents involved in each cross (names given as dam×sire). Northern (N), southern inland (S), and coastal (C) populations were crossed using the crossing design described in text. Bars that share the same lowercase letter are statistically indistinguishable (Tukey’s Honest Significant Differences test, p <0.05).

**Figure 8:**
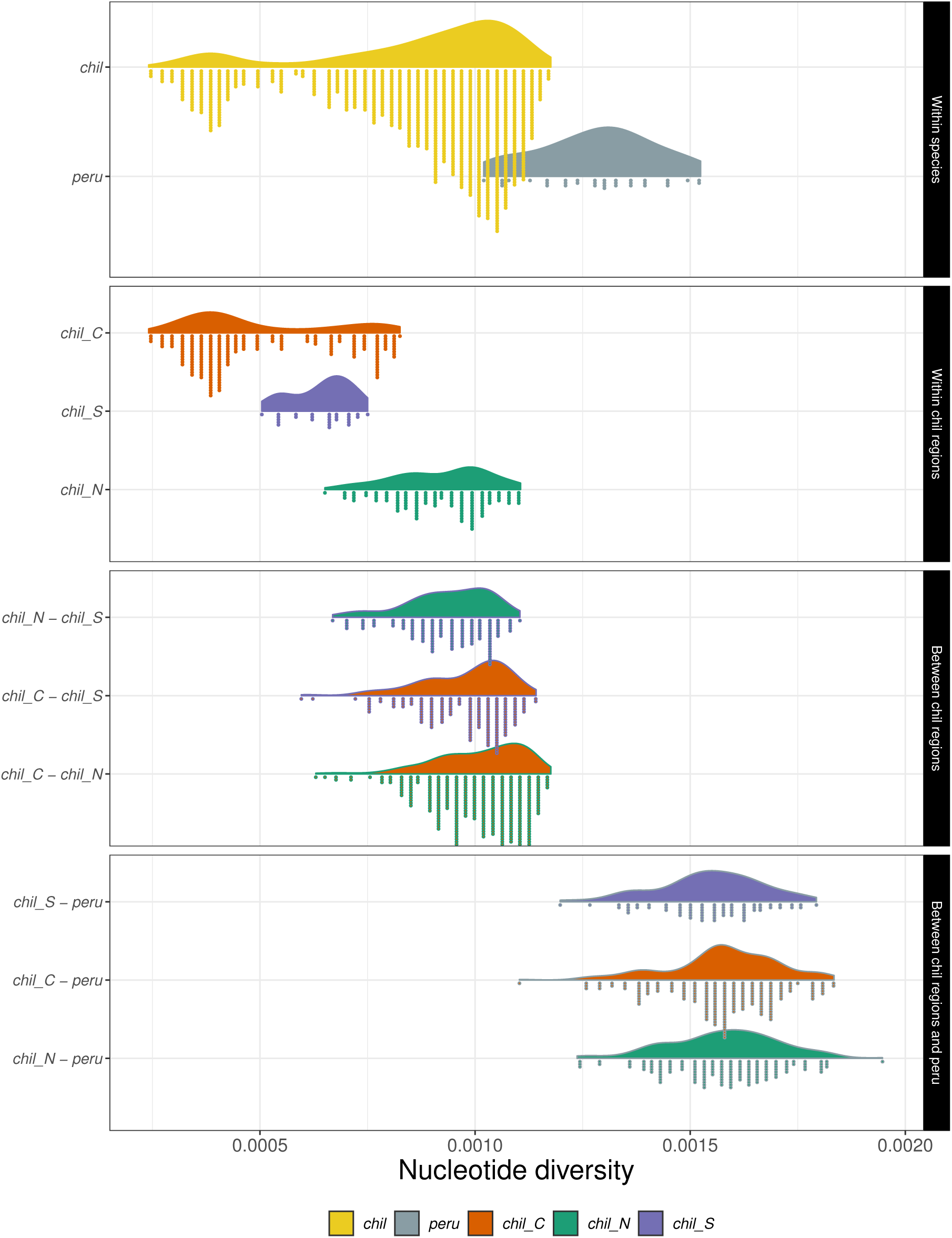
Distribution of nucleotide diversity within (*π*) and between (*D*_*xy*_) populations and species.

**Figure 9:**
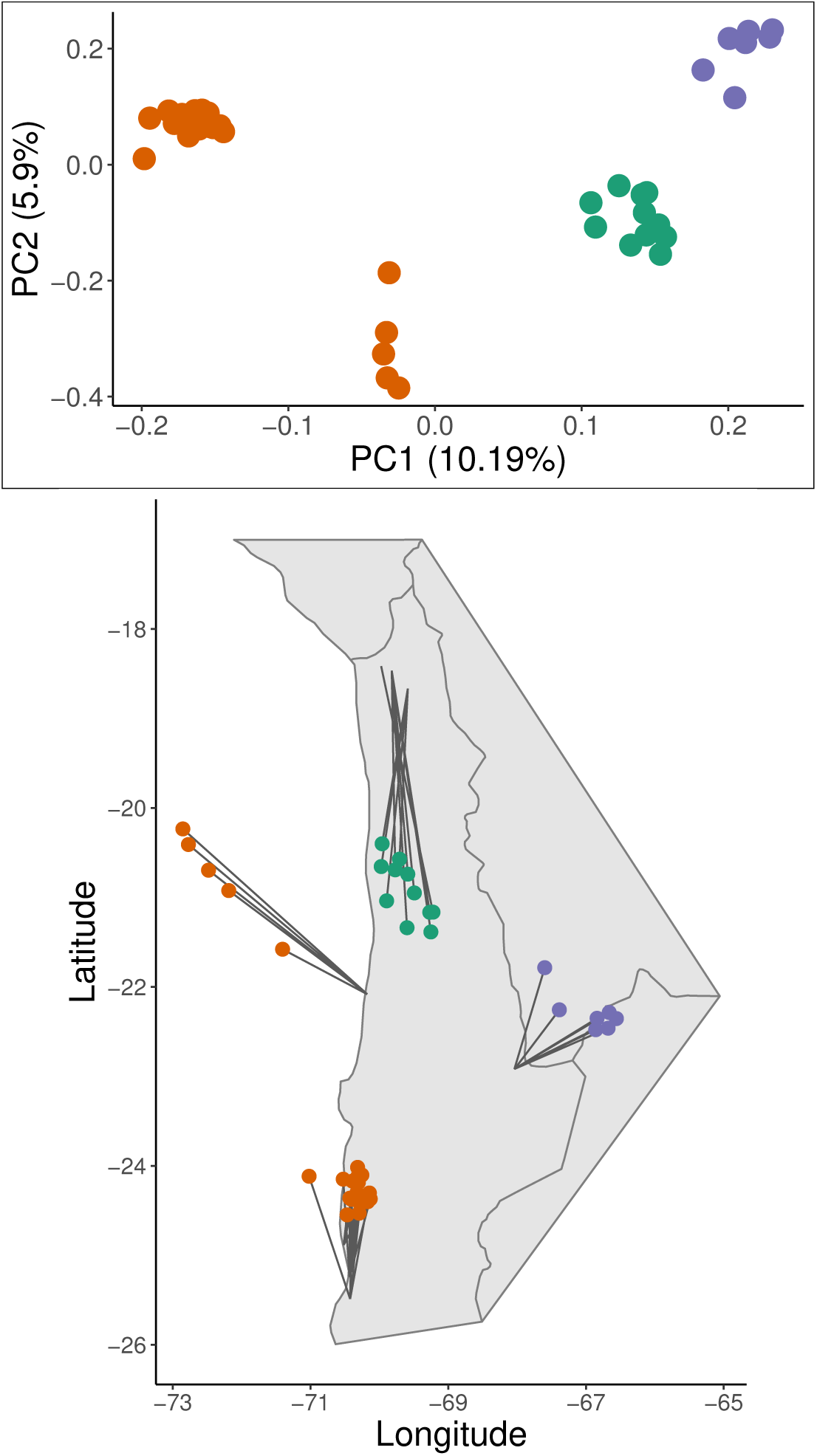
Principal components analysis of *S. chilense* genotypes. Sample PC1 and PC2 coordinates underwent Procrustean transformation onto geographic coordinates to illustrate relationship between genetic variation and geography.

## Discussion

We conducted multiple analyses, each of which provides some weight of evidence that there are at least two separately evolving populations of the currently described *Solanum chilense*. These analyses rely on disparate sources of data and concern multiple aspects of organismal biology, and may provide aid in re-classification of Chilean and Peruvian populations of the *S. chilense* complex. Taken together with previous data, our results suggest that the coastal Chilean populations are geographically, ecologically, morphologically, and genetically distinct from other populations, and could hypothetically experience post-pollination barriers, if they were to re-occur in sympatry. Below, we discuss how they compare to previous studies of variation in the *S. chilense* complex and provide a broader interpretation of the key implications.

### Genetic Differentiation

Phylogenetic analyses reveal a deep divergence between the northern-/western-most populations of *S. chilense* from central Peru and remainder of the complex (Figs. 3, 4). Moreover, the Pacific coast populations, lining lomas formations, from Taltal at the southern margin to Tocopilla at the northern margin, are monophyletic and multiple lines of evidence indicate that they are substantially separated from the rest of the vast interior populations, lining the Andean pre-cordillera. These inland populations span from the area of Salar de Atacama and stretch north to the vicinity of Arequipa (Peru). It is difficult to infer much about the biosystematic relationships involving central Peruvian populations from our data, because they were not included in the our morphological or crossing analyses. Graham (2005), however, performed such crosses and found that these populations form their own crossing group that is at least partially intrinsically reproductively isolated from inland populations. *S. chilense* populations from central Peru are sympatric with both *S. peruvianum, S. corneliomulleri*, and it is unclear whether and how often they experience introgression. Indeed, Bondel et al. (2015) recovered a virtually identical position, as in our analyses, with concatenated nuclear data, and Beddows et al. (2017) found evidence of possible admixture between these sympatric central Peruvian *S. chilense* and *S. peruvianum* (west of the 72nd meridian) in a sampling effort focused on the northern range limit of *S. chilense* and sympatric *S. peruvianum*. They further suggested that central Peruvian *S. chilense* may be a distinct product of recombinational speciation, although the evidence is limited to detection of admixture and somewhat problematic. Specifically, a difficulty faced by most such studies is the certainty of origin of the fidelity between the original collected samples and those used in Beddows et al. (2017); at least one accession (LA0752) may have been mislabelled, out-crossed during seed increases, or otherwise does not correspond to its stated origin (Peru), and appears to originate in Chilean coastal populations (Beddows et al. 2017).

Our RAD-seq data analyses recover a well-supported monophyletic relationship for the coastal population group, and yet it is nested within the rest of the inland species complex members and does not show a particularly clear affinity to either northern or southern inland population group. Previously, Graham (2005) used allozymes, SSRs, and RFLPs to find that the coastal populations may be most closely related to either populations from central Peru, but our analyses do not support this hypothesis. Using sequence data from 30 nuclear loci to estimate relationships among populations, Böndel et al. (2015) found that coastal populations formed a highly supported monophyletic clade that was most closely related to northern inland populations. Our phylogenetic findings are ambivalent regarding the exact placement of coastal populations. RAxML and SVD Quartets phylogenetic analyses reconstructed all coastal plants as monophyletic, however, the two methods differed in the identity of taxa placed sister to the coastal clade (Figs. 3, 4).

Our data support a scenario in which the coastal group is, and has long been, evolving independently, consistent with several other lines of evidence, outlined below. Non-reciprocally monophyletic pattern of relationships for such a species pair (coast vs. inland) is expected under a variety of markers and processes when population sizes are large (Hudson and Coyne 2002), likely to be the case for this obligately outcrossing complex. The level of genetic differentiation, measured by *F*_*st*_, between coastal and southern inland populations is generally higher than the observed level between coastal and northern inland populations, and nearly double the observed differentiation between southern and northern inland populations (Fig. S2). The observed differentiation between the coastal and southern inland populations, which are geographically close, is similar to the interspecific divergence between northern inland *S. chilense* and *S. peruvianum* populations. Furthermore, Structure anal-yses strongly support a cluster of exclusively coastal populations, separated from southern and northern inland samples (Fig. 5). There is no significant support for admixed ancestry between southern inland and coastal populations in our results.

Nucleotide variation is lowest in coastal populations, supporting past studies that have found a similar trend across *S. chilense* populations (Arunyawat et al. 2007; Stadler et al. 2008). Across studied populations, levels of diversity observed in our RAD-seq data are, as expected, lower than previous reports for *S. chilense* (Arunyawat et al. 2007; Stadler *et al*. 2008; Böndel et al. 2015). However, the relative levels of diversity between populations are consistent, and similar. A series of population bottlenecks during a southward range expansion of *S. chilense* is thought to have given way to the observed pattern of reduced nucleotide variation (Stadler et al. 2008; Böndel et al. 2015). The reduced variation observed in the coastal populations may be due to a lower effective population size, increased mating among relatives, or a shift in breeding (and mating) system away from self-incompatibility. Our sampled coastal populations are the southern range limit of all wild tomato species and are the result of range expansion from the north, the center of wild tomato diversity (Rick and Lamm 1955; Peralta et al. 2008; Böndel et al. 2015). Peripheral plant populations are often associated with self-compatibility and therefore may experience reduced effective population sizes due to the increased potential for self-fertilization (Baker 1955, 1967; Stebbins 1957; Herlihy and Eckert 2005; Pannell et al. 2015). Ongoing work in our lab has uncovered some variation in breeding system, indicating that self-compatible mutations may be segregating in coastal Chilean populations (ARR and BI, pers. obs.).

We sequenced two sets of alleles, SX-30 and S3-6, from coastal and both inland populations, which display variation within functionally identical allelic classes, consistent with considerable divergence between the coastal populations and inland populations (Fig. 2, Igic et al. 2007). The rationale for this claim is as follows. Due to strong negative frequency-dependent selection, dozens of S-alleles can be maintained within species (Lawrence 2000). Loss and fixation of alleles by drift is strongly countered by this mode of selection, alleles persist for very long periods of time (Clark and Kao 1994). Takahata (1990) showed that the between-allele and within-allele (same functional type with neutral sequence differences) coalescent times can be studied with mathematical tools developed for neutral loci, except that they must be scaled to account for changes in relative effective population size at these two hierarchical levels of S-allele evolution. S-alleles may be extremely sensitive markers for population structure and gene flow, because rare migrant alleles are “vacuumed” into populations that lack them by negative frequency-dependent selection. It aids immigrant S-alleles in a population, as rare alleles are favored by selection and less likely to be lost due to drift (Wright 1939; Vekemans and Slatkin 1994; Schierup et al. 2000). Strong empirical support for this conjecture is broadly lacking (Muirhead 2001; Glémin et al. 2005; Joly and Schoen 2011), mainly on account of difficulty in collecting of appropriate data—capturing and sequencing identically functional, but sequence-variable alleles. Nevertheless, studies have shown that within-allele genealogies reflect species phylogeny and can, in principle, detect structure (Lu 2001; Castric et al. 2008). While various unique features of evolution of S-alleles have been used to infer demographic changes from distant past (Paape et al. 2008; Busch et al. 2011) and to reconstruct ancestral states of plant breeding systems (Igić et al. 2008), patterns of variation within S-allele types have not been used extensively for detection of population structure, despite the exciting opportunities to do so (Vekemans and Slatkin 1994).

### Morphological differentiation

Morphological characterization of this taxon is important in part because nearly all historical taxonomic treatments exclusively used herbarium specimens. Herbarium specimens of the type specimen used by Philippi (1860) to describe what would later become *S*. chilense were collected from coastal Chile (then Bolivia). Muller 1940 noted that without further context, the placement of coastal and inland populations into the same taxon based on morphology was tenuous, because of the amount of variation across *S. chilense*’s range. Rick and Lamm (1955) did not examine the coastal collections of *S. chilense* for their morphological analyses, but a single herbarium specimen from the coastal localities they examined resembled *S. chilense* rather than *S. peruvianum*. Peralta et al. (2008) included the coastal specimens from other *S. chilense*.

Our direct measurement of morphological traits from plants in the field are most thorough investigation of morphological diversity of this group, in terms of the number of sampled individuals and populations. Glasshouse-grown plants strongly resemble plants measured in the field from their respective groups, indicating that the differences observed in the field are not entirely environmentally dependent (ARR, unpub. data). Despite extensive morphological variation within all populations of this self-incompatible taxon, coastal populations are easily identifiable in the field (and glasshouse) by their unusually low prostrate growth, a trait that we did not measure or include in our study. They also occupy a fairly distinct space in the PCA analyses, although they partly overlap. And yet, *S. chilense* and *S. peruvianum*, two similar, widely recognized species (Rick and Lamm 1955; Peralta et al. 2008) also overlap in the same analyses. Our field experience and previous work (Peralta et al. 2008; Chetelat et al. 2009) indicates that closely related wild tomato species may be difficult to delimit with morphological diagnostic characters, because of both high intraspecific variation and low interspecific divergence.

### Post-pollination barriers within Solanum chilense (Dunal) Reiche

We examined a small sliver of total reproductive isolation by estimating barriers to seed set from inter-population crosses, following manual cross-pollination. Crosses involving coastal plants as either a female or male parent resulted in significantly fewer seeds per fruit than all other crosses (Fig. 7). Graham (2005) also found significant reductions in seed set per fruit for inter-population crosses involving coastal and inland accessions. Levels of interspecific reproductive isolation between *S. chilense, S. peruvianum*, and their close relatives, measured as hybrid seed formation, vary from nearly complete to non-existent (Rick and Lamm 1955; Beddows et al. 2017; Roth et al. 2017). Although the observed reduction in seed set from coastal and inland crosses is not absolute, it is comparable to interspecific crosses between *S. chilense* and *S. peruvianum* (Rick and Lamm 1955; Beddows et al. 2017). The coastal population group is of specific interest, because it is genetically distinct and notably ecogeographically isolated, separated from the rest of the taxon by some of the driest land on the planet, and occurs at lower altitudes (Fig. 1). Thus early-acting, pre-mating barriers likely contributed more to the recent and ongoing reproductive isolation between the coastal and inland populations of *S. chilense*. The closest southern inland populations are separated from the coastal ones by ca. 100-200km of utterly barren desert and approximately 2400-3600m in elevation.

We do not describe a new taxon in this paper, but we believe that a practical approach to taxonomy strongly justifies doing so. One of the major lines of work in our lab concerns estimation of alternate breeding system effects on rates of speciation, extinction, and transition (Goldberg et al. 2010). Clearly, such inference can be critically affected subject by species delimitation. For instance, if a monograph were to rely on delimitation criteria that systematically over-estimate numbers of self-fertile species and underestimate numbers of outcrossing species, the resulting inference would be confounded—it could plausibly recover higher speciation (and maybe extinction) rates of self-fertile species.

Although species in *Solanum* sect. *Lycopersicon* have undergone a number of recent taxonomic changes (Peralta et al. 2008), self-fertilizing species have enjoyed relative stability compared with their self-incompatible cousins. Indeed, taken together Böndel et al. (2015); Pease et al. (2016); Beddows et al. (2017), as well as the present study, each hint that the taxonomic status of nearly all obligately outcrossing species remains unsettled. These species all likely have enormous effective population sizes, result in correspondingly greater coalescent times, which yield unfavorable conditions for recovery of reciprocal monophyly at any marker, whether genetic or morphological. This makes it difficult to easily uncover fixed eye-catching morphological differences, useful for clean binary key construction. It therefore seems prudent to closely examine the problem of species delimitation in outcrossing groups, such as the *S. chilense* complex, as well as consider the wider consequences of their under-description. Our study indicates a strong potential for taxonomic studies to systematically lump obligately outcrossing species, a practice that is certain to cause a bias in estimates of speciation, extinction, and transition rate estimates for breeding system traits and their correlates—a serious problem for already suspect macroevolutionary models used across many fields of biology (Louca and Pennell 2020).

### Conclusions

Overall, each of our analyses provides some weight of evidence that the Pacific coastal populations and Andean inland populations of the currently described *Solanum chilense* (Dun.) Reiche represent separately evolving populations. A variety of species concepts agree that the defining property of species are separately evolving populations, although they often adopt different properties acquired by lineages during the course of (sufficient) divergence (De Queiroz 2007). Populations in the *S. chilense* complex show substantial genetic and morphological differentiation, they are completely allopatric, and show post-pollination reproductive barriers, hinting that continued separation is plausible in a hypothetical return of sympatry. Moreover, they differ, at least anecdotally, in key ecological traits, including edaphic and altitudinal suitability. Taken together with previous data, our findings seem to strongly support that minimally two separate species, under a number of concepts, comprise the *S. chilense* complex. Finally, the northern-most populations, from central Peru, may also form a separate species, although the apparent hybridization with *S. peruvianum* (Beddows et al. 2017), and lack of extensive collections and data on this disjunct group complicates its status.

*S. chilense* also provides a unique opportunity for an examination of the genetic basis, tempo and mode of accumulation of a variety of isolating barriers (Moyle 2008) between obligately outcrossing populations and species, which often maintain astonishingly large long-term effective population sizes. Transplant experiments measuring inland and coastal hybrid viability and fertility in the field are warranted, as is detailed genetic work, needed to investigate the demographic history of this complex. Moreover, this taxon is broadly sympatric with *S. peruvianum*, and presents intriguing possibilities for further work on detection of any patterns of introgression (Durand et al. 2011). Further studies detailing the components of reproductive isolation (Ramsey et al. 2003) are especially promising in this group. Our key findings also show that, despite its vast economic importance, *Solanum* sect. *Lycopersicon* still exhibits considerable taxonomic instability.

## Acknowledgements

We are grateful to Schaal lab at Washington University in St. Louis, and specifically W. A. Smith, for field collections of 1993/1994, which turned our attention to the tomato group and provided invaluable preliminary data. Kelly Robertson performed many of the greenhouse crosses. Jim Folsom and Matt Ritter successfully advocated for the ‘old-school’ biosystematic approaches and inspired the our progress on this paper. Phillip Fenberg helped perform measurements and fixed the cable. This work was supported by the National Science Foundation grants NSF DEB-0919089 and DEB-1655692. Computational resources were provided by the Research Computing group at the University of Illinois at Chicago.

---

**Table S1:**
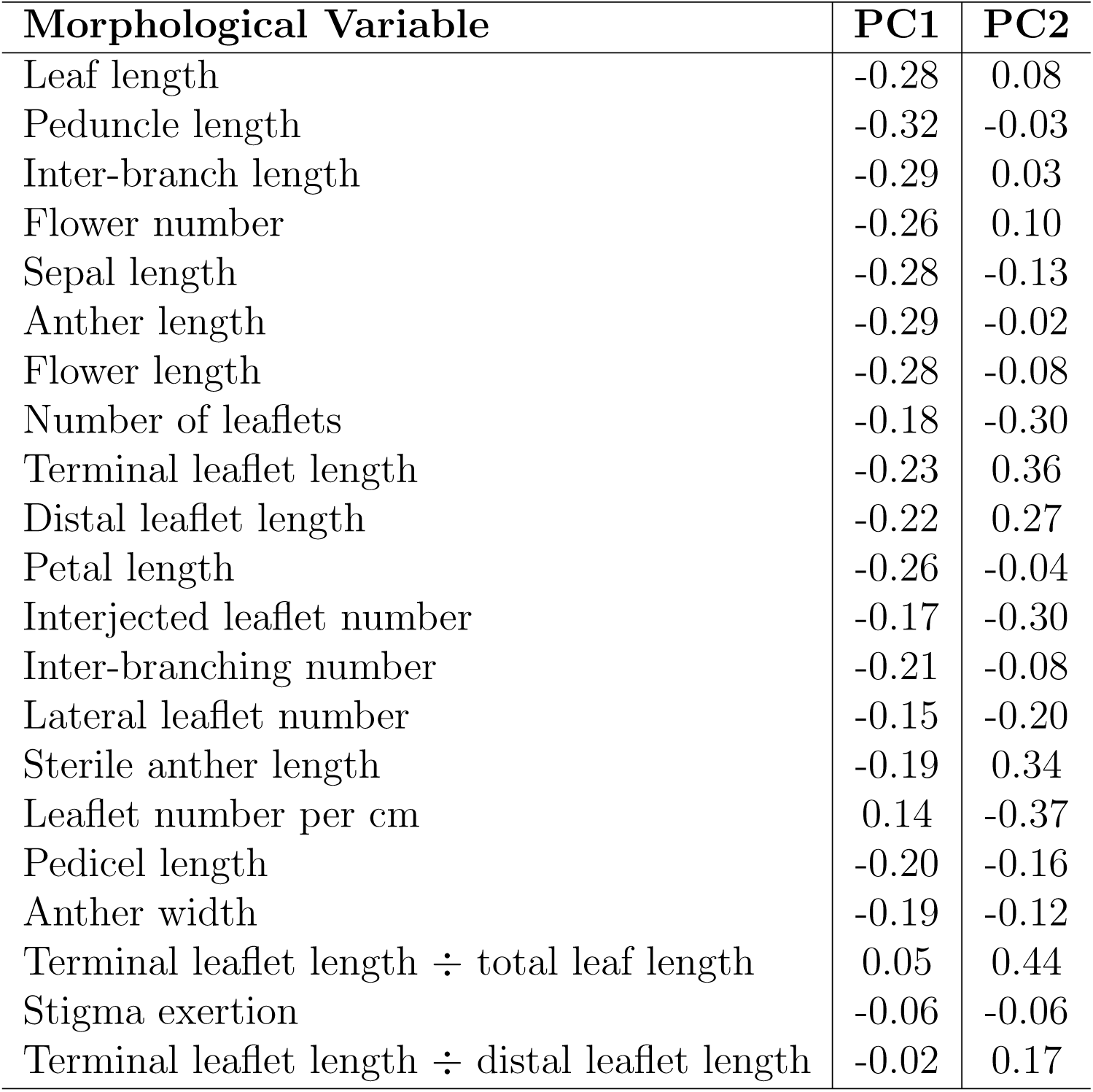
The loadings of each morphological measurement on the first two principal components. Leaf and inflorescence size measurements contributed the most to the first principal component, while leaflet size and leaf shape measurements contributed the most to the second.

| **Morphological Variable** | **PC1** | **PC2** |
| --- | --- | --- |
| Leaf length | -0.28 | 0.08 |
| Peduncle length | -0.32 | -0.03 |
| Inter-branch length | -0.29 | 0.03 |
| Flower number | -0.26 | 0.10 |
| Sepal length | -0.28 | -0.13 |
| Anther length | -0.29 | -0.02 |
| Flower length | -0.28 | -0.08 |
| Number of leaflets | -0.18 | -0.30 |
| Terminal leaflet length | -0.23 | 0.36 |
| Distal leaflet length | -0.22 | 0.27 |
| Petal length | -0.26 | -0.04 |
| Interjected leaflet number | -0.17 | -0.30 |
| Inter-branching number | -0.21 | -0.08 |
| Lateral leaflet number | -0.15 | -0.20 |
| Sterile anther length | -0.19 | 0.34 |
| Leaflet number per cm | 0.14 | -0.37 |
| Pedicle length | -0.20 | -0.16 |
| Anther width | -0.19 | -0.12 |
| Terminal leaflet length ÷ total leaf length | 0.05 | 0.44 |
| Stigma exertion | -0.06 | -0.06 |
| Terminal leaflet length ÷ distal leaflet length | -0.02 | 0.17 |

**Table S2:**
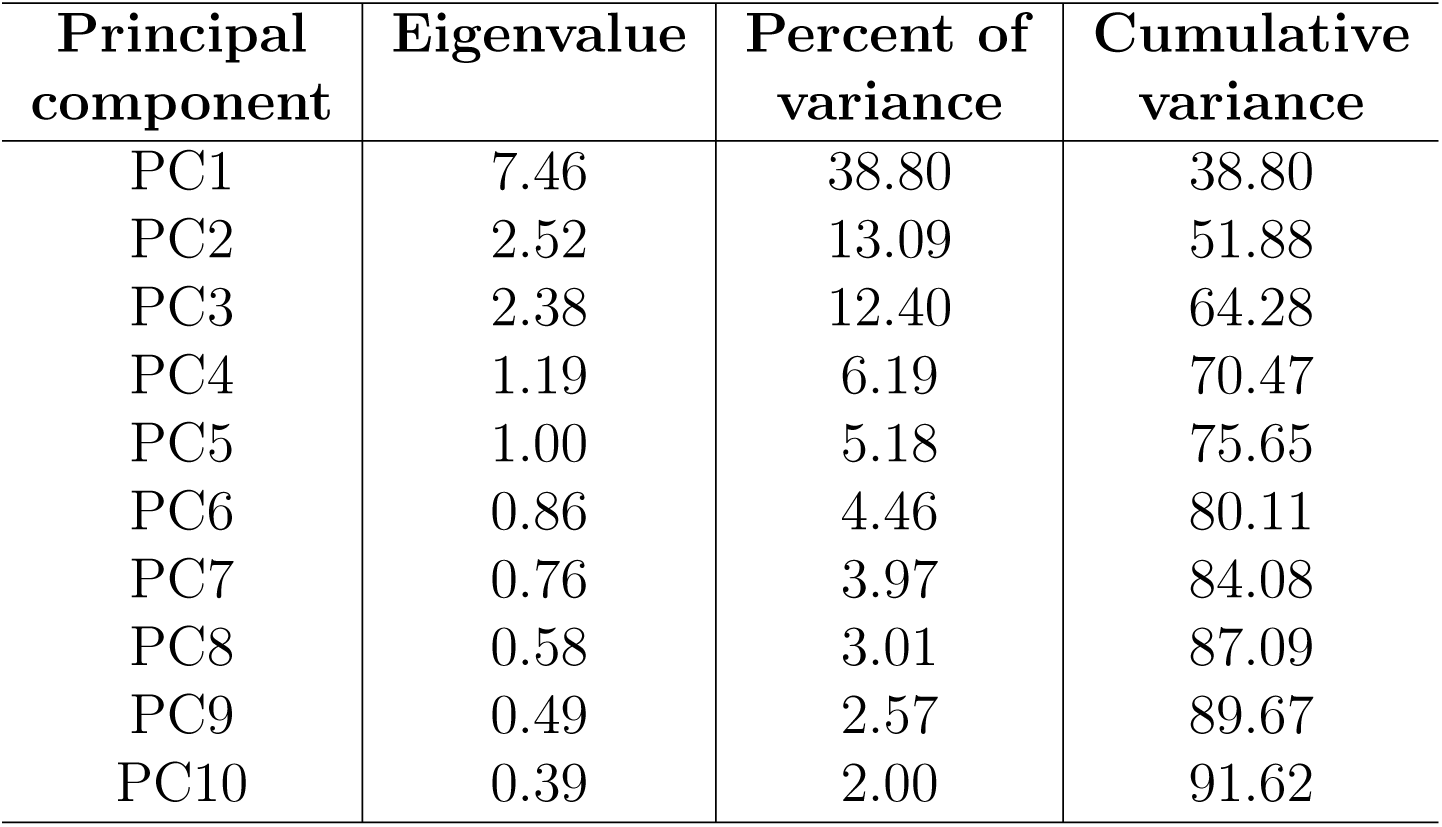
Eigenvalues and the amount of variation of the morphological dataset explained by the first ten principal components.

**Table S3:**
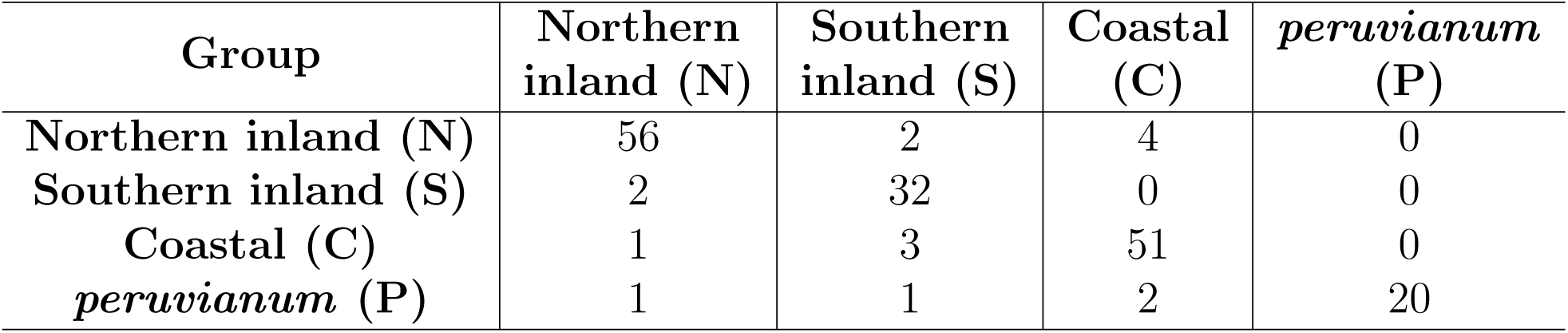
Placement of individuals into predicted groups by linear discriminant analysis (LDA), using the collected morphological floral and vegetative data. Correct group memberships are given in each row. LDA predicted membership assignments are given in each column.

**Figure S1:**
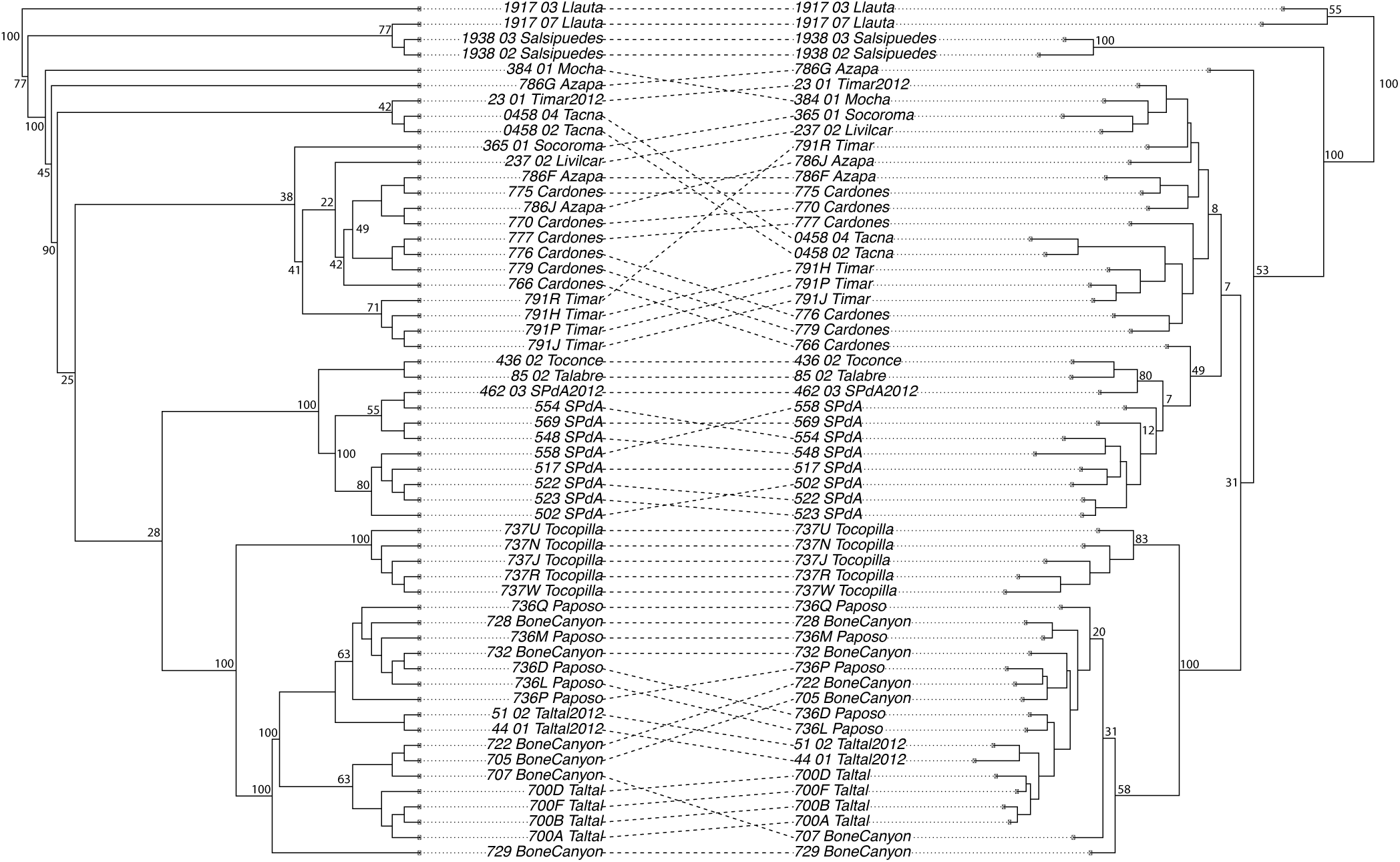
Co-plotted SVD Quartets (left) and RAxML (right) phylogenies comparing reconstructed topologies. One major disagreement between the two phylogeny reconstruction methods is the identity of taxa that are most closely related to the coastal *S*. chilense clade. With low bootstrap support (28%), SVD Quartets places a highly supported southern inland clade (100%) as sister to the coastal clade. RAxML places the coastal *S*. chilense clade as sister to a larger paraphyletic clade consisting of northern and southern inland individuals.

**Figure S2:**
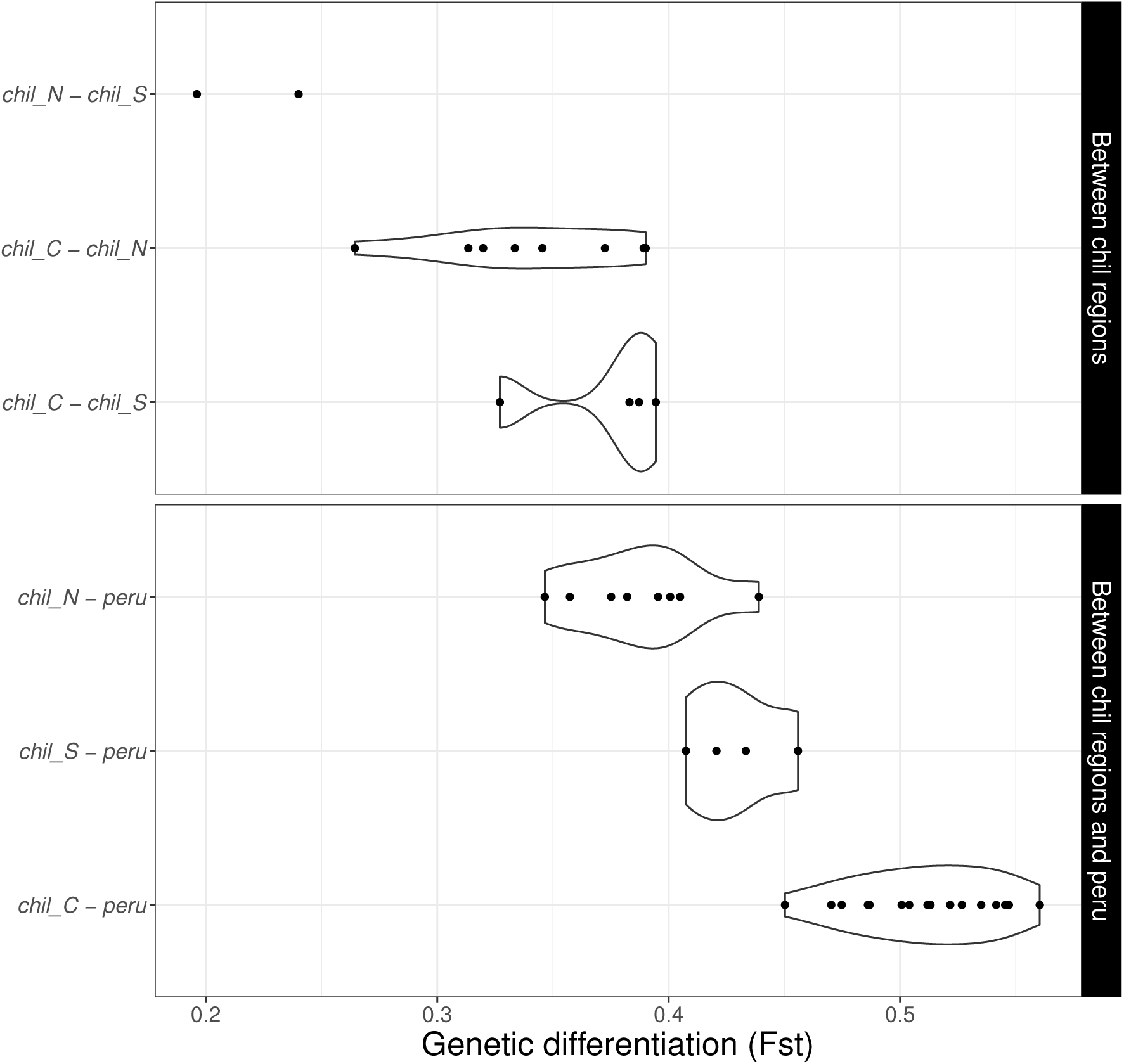
Estimates of pairwise genetic differentiation (*F*_*st*_) between populations of *S. chilense* and *S. peruvianum*. Comparisons are separated by within *S. chilense* and between groups (top panel), and interspecifically between *S. chilense* groups and *S. peruvianum* (bottom panel).

**Figure S3:**
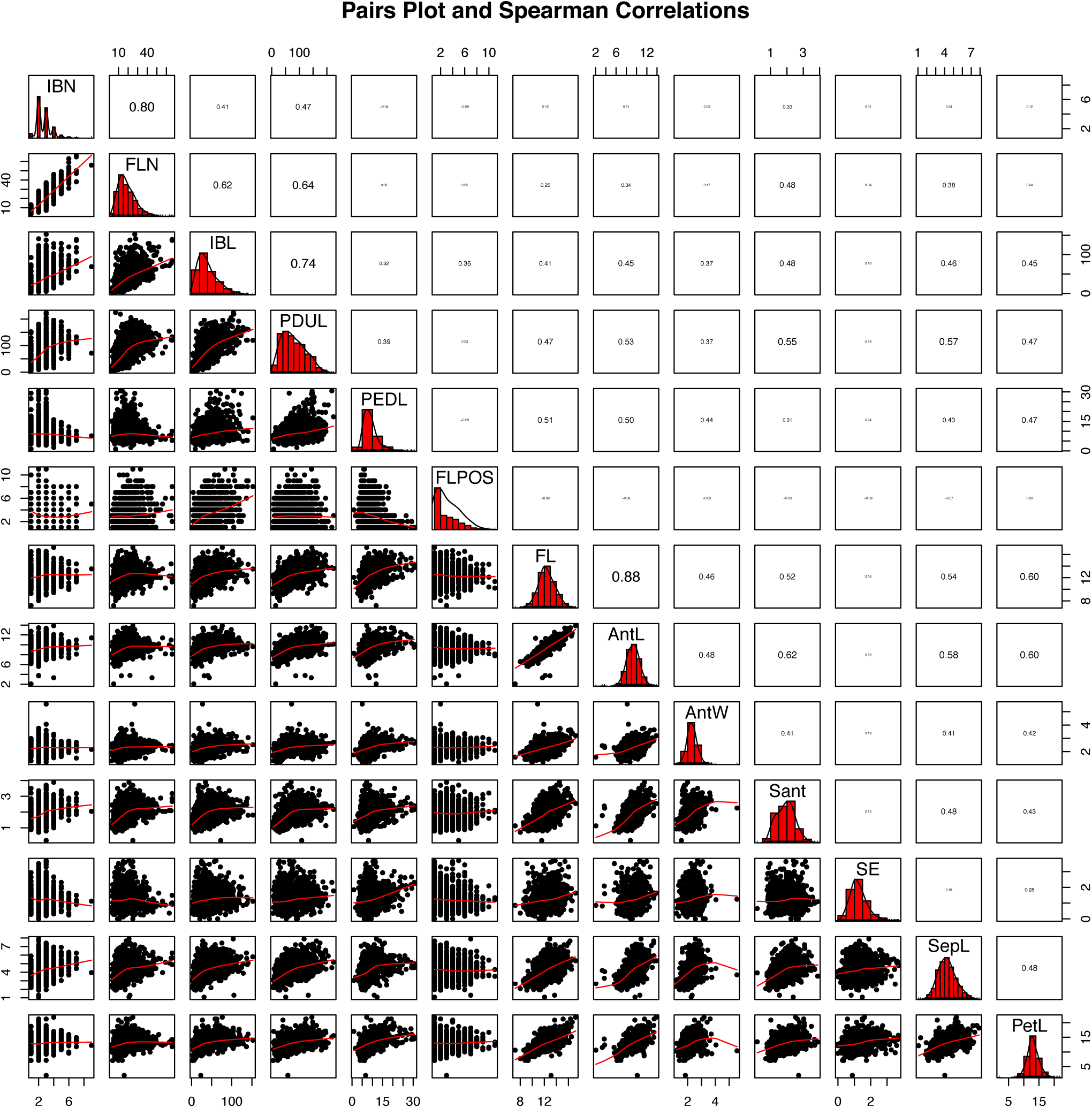
Pairs plots, distributions, and Spearman correlations of measured morphological traits. The bottom triangle summarize bivariate plots, diagonal plots summarize variation, and top triangles show Spearman correlations. Legend: IBN=inter-branching number, FLN=flower number, IBL=inter-branching length, PDUL=peduncle length, PEDL=pedicel length, FL=flower length, AntL=anther length, AntW=anther width, Sant=sterile anther length, SE=stigma exertion, SepL=sepal length, and PetL=petal length.

